# RBPMS promotes contractile smooth muscle splicing and alters phenotypic behaviour of human embryonic stem cell derived vascular smooth muscle cells

**DOI:** 10.1101/2022.11.27.516868

**Authors:** Aishwarya G Jacob, Ilias Moutsopoulous, Alex Petchey, Irina Mohorianu, Sanjay Sinha, Christopher WJ Smith

## Abstract

Differentiated Vascular Smooth Muscle Cells (VSMCs) express a unique network of splice isoforms (smooth muscle specific alternative splicing - SM-AS) in functionally critical genes including those comprising the contractile machinery. We previously described RNA Binding Protein Multiple Splicing (RBPMS) as a potent driver of contractile, aortic tissue like SM-AS in VSMCs using rodent models. What is unknown is how RBPMS affects VSMC phenotype and behaviour. Here, we use human embryonic stem cell-derived VSMCs (hES-VSMCs) to dissect the role of RBPMS in SM-AS in human cells and determine the impact on VSMC phenotypic properties. hES-VSMCs are inherently immature and display only partially differentiated SM-AS patterns while RBPMS levels are undetectable endogenously. Hence, we used an over-expression system and found that RBPMS induces SM-AS patterns in hES-VSMCs akin to the contractile tissue VSMC splicing patterns in multiple events. We present *in silico* and experimental findings that support RBPMS’ splicing activity as mediated through direct binding and via functional cooperativity with splicing factor RBFOX2 on a significant subset of targets. Finally, we demonstrate that RBPMS is capable of altering the motility and the proliferative properties of hES-VSMCs to mimic a more differentiated state. Overall, this study emphasizes a critical splicing regulatory role for RBPMS in human VSMCs and provides evidence of phenotypic modulation by RBPMS.

## Introduction

Vascular smooth muscle cells (VSMCs) constitute the major architectural component of the large arteries and in normal healthy vessels, present in a mature or differentiated, contractile phenotypic state. This is reflected in their function as effectors of vascular tone and mediators of tonic contraction to ensure effective blood flow (vasoconstriction and vasodilation). The molecular profiles of these cells are reflective of these properties and several markers including the expression of smooth muscle myosin heavy chain (SM-MHC or MYH11), Calponin1 (CNN1) and TAGLN are associated with and are functionally relevant for the contractile state. Notably, VSMCs are phenotypically plastic and in response to vessel wall injury and in several cardiovascular conditions including atherosclerosis, restenosis and hypertension, they dedifferentiate into more mesenchymal states that are associated with highly proliferative and synthetic characteristics. This switch is accompanied by major changes in the cellular transcriptome including loss of expression of the contractile marker network. More recently, with the advent of single cell RNA sequencing, we have been able to dissect these molecular changes in far greater detail and better appreciate the phenotype switching process and the heterogeneity underlying contractile VSMCs of different embryonic lineages [1-5]. However, most of our knowledge of the molecular networks characterising these VSMC states is centred around the mRNA abundance profiles of marker genes, leaving us, currently, with a view only of the transcriptional component of the VSMC transcriptome.

Alternative pre-mRNA splicing (AS) is a tightly regulated, pervasive post-transcriptional process that facilitates the generation of multiple, often functionally distinct transcript isoforms from the same gene. Previously, we showed that AS is a major component that defines arterial tissue VSMC transcriptomes. We uncovered a network of unique mRNA isoforms expressed in contractile VSMCs generated as a result of Smooth Muscle specific Alternative pre-mRNA splicing (SM-AS) [6, 7]. This includes well known markers of advanced VSMC maturity such as heavy-Caldesmon (CALD1) [8] and meta-Vinculin (VCL) [9, 10] and the inclusion of smooth muscle specific mutually exclusive exons in genes like *TPM1* and *ACTN1* which allows for distinction from the isoforms expressed in cardiac and skeletal muscle, and non-muscle cells. Functionally, the genes affected by SM-AS largely pertain to smooth muscle contractility, the acto-myosin cytoskeletal network and cell adhesion, although, it is not clear in all the transcripts how SM-AS ultimately alters protein domain and activity. SM-AS is lost during phenotype switching alongside the contractile marker network expression [6, 7].

The regulation of AS is an intricate and complex process that is tightly connected to the expression, localisation and activity of a large class of proteins called RNA Binding Proteins (RBPs). Usually, with one or more RNA Recognition Motifs (RRMs) or other RNA binding domains, these proteins bind target pre-mRNAs with varying affinities at defined or more degenerate motifs and, in concert with each other, direct splicing activity to specific sites. This finely controls the level of AS isoforms being expressed and is an important attribute of cell-type or tissue specification [11]. Numerous RBPs including QKI and members of the CELF and MBNL protein families have been described with roles in the cardiovascular system – both developmentally and in the context of disease, where they perform a number of roles including splicing control [12-18]. In VSMCs, a handful of RBPs including QKI [19], HuR [20], hnRNPA2B1 [21] and PTBP1 [6] have been described as targeting key regulators of VSMC phenotype and transcription in the context of phenotype switching and diseases like hypertension [22]. Another family of critical AS regulators comprises the RBFOX genes. Primarily described with roles in neuronal and striated muscle developmental programmes including cardiac muscle [23-31], members of this family, especially RBFOX2, are heavily associated with heart failure conditions [18, 32-37]. In the vasculature, a splicing regulatory role for RBFOX2 has been described in the endothelium [38] and in VSMCs, which express high levels of this protein, for only a few events including calcium channel genes (*CACNA1C*) [39].

The RNA Binding Protein Multiple Splicing (RBPMS) family of genes - RBPMS and RBPMS2 - are now being increasingly appreciated as critical regulators of smooth muscle and cardiac post-transcriptional RNA processing with developmental roles [40-42]. These proteins were originally identified in the developing heart as RBPs [43, 44] and RBPMS is expressed in the murine myocardium from E12.5 and in vasculature from E15.5 [45]. RBPMS2 was highlighted with functions pertaining to visceral smooth muscle [46]. In vascular smooth muscle, RBPMS is super-enhancer associated and highly expressed in mature, aorta tissue VSMCs and on this basis was proposed as a potential master regulator of SM-AS [7]. Consistent with this, like MYH11 and other VSMC identity markers, RBPMS is a part of a contractile gene network identified by single cell transcriptome profiling [2] and its expression is dynamically regulated during phenotype switching [7]. A recent study definitively showed RBPMS as an essential gene with homozygous deletion resulting in neo-natal lethality. Its functional importance and the resulting lethality were attributed to under-developed myocardium although some vascular phenotypes were observed [45]. The role of RBPMS in vascular function and VSMC phenotype as a splicing regulator remains unclear. We recently described RBPMS, as a conserved, critical regulator of SM-AS. Overexpression of RBPMS in phenotypically switched rat PAC1 VSMCs was sufficient to promote tissue-VSMC like splicing patterns. Moreover, increased RBPMS expression fully accounted for nearly 20% of the SM-AS network regulated between partially differentiated and undifferentiated PAC1 cells. Based on these, and other criteria [47], we proposed that RBPMS acts as a master splicing factor in VSMCs [7].

Here, we hypothesized that RBPMS plays a critical role in mature VSMCs and the SM-AS network underlies differentiated or contractile VSMC phenotypic behaviour. We test this using the human embryonic stem cell derived VSMC (hES-VSMCs) model [1]. Although these cells are *bona fide* VSMCs, we found that they express basal level splicing patterns and no detectable endogenous RBPMS, consistent with a fetal/immature VSMC phenotype. We show using over-expression strategies, that RBPMS is a potent driver of SM-AS akin to tissue VSMCs modulating the actin-cytoskeleton, focal adhesion and contractile machinery. We show that RBPMS functionally interacts with and requires RBFOX2, at a subset of these targets to mediate SM-AS. Finally, we describe how RBPMS over-expressing hES-VSMCs show lowered motility and proliferation in a manner similar to mature, differentiated VSMCs. Overall, we make a case for the master splicing regulator RBPMS as a driver of phenotypic behaviours associated with mature, contractile tissue VSMCs.

## Results

### Human Embryonic Stem cell derived Vascular Smooth Muscle cells (hES-VSMCs) do not endogenously express detectable levels of RBPMS protein

To determine the impact of RBPMS in human VSMCs, we decided to use the human embryonic stem cell (ESC) derived model. Briefly, human ESCs were differentiated into neural crest (NC) or lateral plate mesoderm (LM) intermediate lineages using established protocols [1, 48, 49]. Following this, the cells were subjected to PDGF and TGFβ treatment in basal media for 12 days[1, 48, 49] and then maintained in 10% FBS containing VSMC-media to mature hES-VSMCs. Compared to ESCs and HEK293T cells, hES-VSMCs from both lineages not only display increased levels of several VSMC markers [1, 48, 49] but also increased SM-AS patterns. In some genes like *ITGA7* [6] there is a complete switch to the SM-AS isoform similar to differentiated or mature tissue VSMCs (Fig 1A, Fig 1 associated supplementary 2A). But for others such as *MYOCD* [7] the switch in splicing is only partial, with lower levels of exon inclusion than are seen in adult arterial tissue (Fig 1A, Fig 1 associated supplementary 2B). However, when we examined the hES-VSMCs for RBPMS expression, we could not detect endogenous protein although base-line level of *RBPMS* transcripts were detectable by quantitative RT-PCR (Fig 1 associated supplementary 1B). Rather, the levels of RBPMS decreased during differentiation from embryonic stem cells (H9 ESCs) to hES-VSMCs via both the NC and LM lineages with the intermediate stages showing variable levels of the protein (Fig 1 associated supplementary 1C). Because RBPMS is highly co-expressed with advanced VSMC marker MYH11 in arterial VSMCs (Fig 1 associated supplementary 1A – human heart cell atlas [5]), it is likely that RBPMS is also only highly expressed in mature adult VSMCs and far less so in the more immature or foetal-like hES-VSMCs.

**Fig 1.**
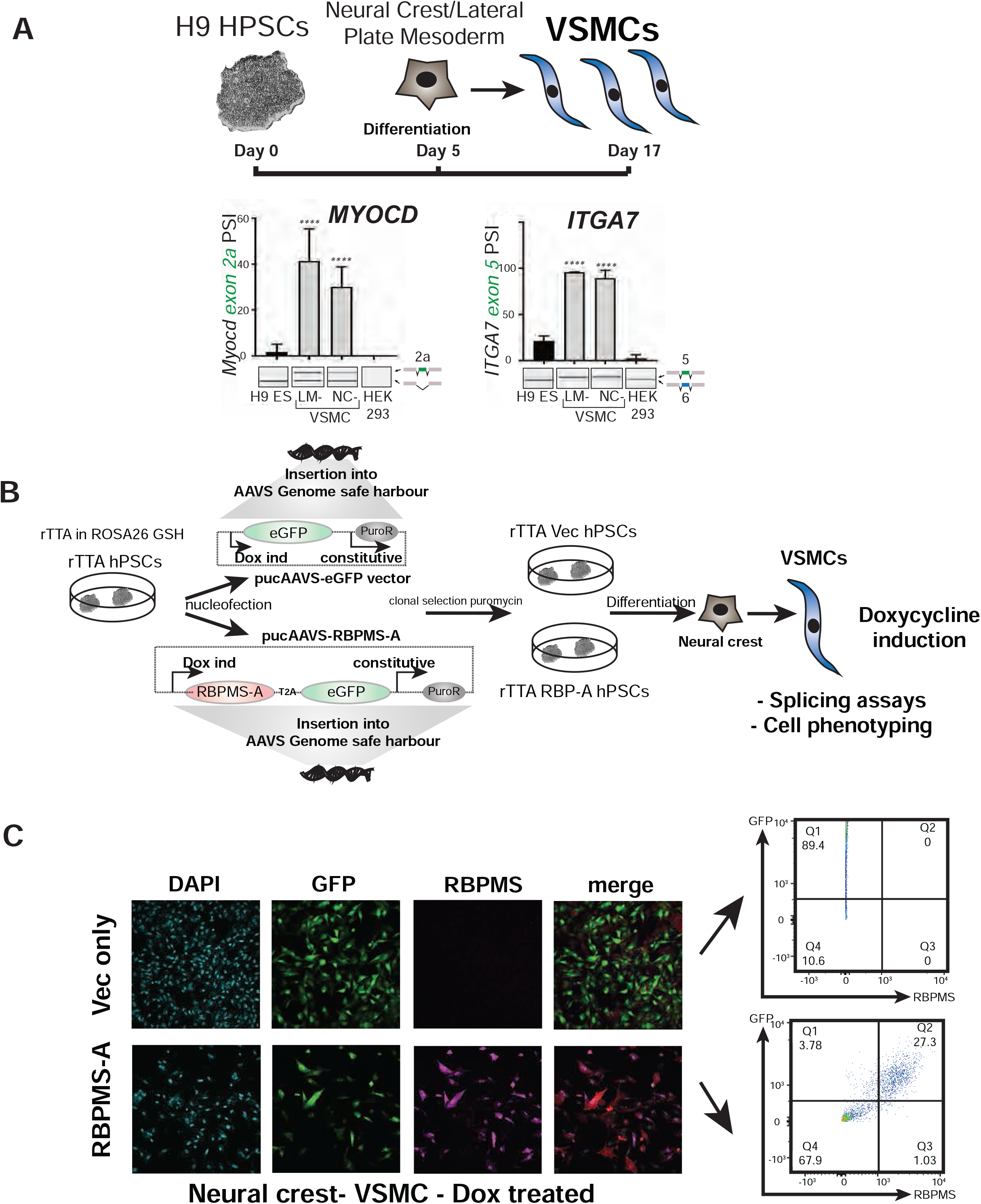
hES-VSMC model to investigate RBPMS role in SM-AS. **A**. H9-human embryonic stem cells were differentiated into VSMCs via the neural crest or the lateral plate mesoderm lineage. RT-PCR of splicing targets *MYOCD* and *ITGA7* are shown. Graphs show <u>P</u>ercent <u>S</u>pliced In (PSI) of the SM-specific exons (green) using data from the electrophoretic images below. N=3. **B**. Schematic showing the H9-rTTA system. H9-rTTA embryonic stem cells heterozygous for the rTTA protein in the ROSA26 locus were nucleofected with pUC-AAVS-eGFP vectors containing either GFP (rTTA Vec hESCs) or RBPMS-T2A-GFP (rTTA RBPMS-A ESCs) under the control of Doxycycline inducible promoters. Selected clones of both ESC lines were differentiated via the neural crest lineage into VSMCs, induced with 0.2ug/ml Doxycycline in 10% FBS containing VSMC media for at least 5 days before use for splicing and phenotyping assays. **C**. Representative Immunofluorescence images showing rTTA Vec and rTTA RBPMS-A ES derived VSMCs via the neural crest lineage (N=3) with mosaic GFP expression correlating with detectable levels of RBPMS protein only in the RBPMS-A line. Representative validation by flow cytometry staining for RBPMS with only the GFP positive RBPMS-A VSMCs expressing the over-expressed RBPMS (N=3).

### Doxycycline-inducible over-expression in hES-VSMCs reveals RBPMS as a major driver of modulations in the transcriptome

We hypothesized that the immature AS patterns in hES-VSMCs might be due to the lack of RBPMS expression. To address this, we generated an inducible over-expression platform. We inserted constructs bearing GFP alone (Vector/Vec lines) or RBPMS linked to GFP via a T2A sequence (facilitates independent translation of RBPMS and GFP from a single poly-cistronic transcript) (RBPMS-A/RBP lines) under the control of Doxycycline –inducible promoters into the pUC-AAVS genomic safe harbour of the H9-rTTA ESC line[50]. This hESC line is heterozygous at the ROSA26 locus for the reverse transactivator gene (rTTA) and constitutively expresses the rTTA protein. Once puromycin-resistant clones were isolated, the VEC and RBPMS hESC lines were expanded and differentiated via the LM or the NC intermediates into hES-VSMCs. Doxycycline (0.2ug/ml) was added to the media and cells were treated for at least 5 days before expansion and use for molecular and phenotypic analyses (Fig 1B).

We observed that our Dox-inducible over-expression system was inherently mosaic with cells within the same population expressing varying levels of GFP and corresponding levels of RBPMS (Fig 1C). This variation was seen in different clones of RBPMS inducible cells. This is likely due to partial and stochastic epigenetic silencing at the pUC-AAVS locus as sodium butyrate treatment (pan HDAC inhibition) allowed for GFP expression in all Doxycycline inducible cells (data not shown). With this pilot observation we had the confidence to proceed with these cells and used this mosaicism to our advantage as an internal control to compare Doxycycline treated cells that express low and high levels of RBPMS under identical culture conditions. We used FACS to sort the doxycycline-treated RBPMS-hES-VSMCs based on their GFP intensity, which correlated with RBPMS expression (Figs 1C, 2A), and isolated RNA from the various fractions. In addition, as a control, we used the vector lines that expressed GFP alone upon Doxycycline induction and sorted these cells based on GFP intensity. RT-PCR for known targets of RBPMS-induced splicing [7] showed that only high GFP i.e. high RBPMS expressing cells from the RBPMS-hES-VSMCs displayed tissue-like contractile SM-AS. The corresponding low GFP RBPMS-hES-VSMCs populations and both high and low GFP samples from the vector hES-VSMCs displayed baseline AS (Fig 2B). This was verified in VSMCs from both the LM and the NC lineages (Fig 2B and Fig 2 associated supplementary).

**Fig 2.**
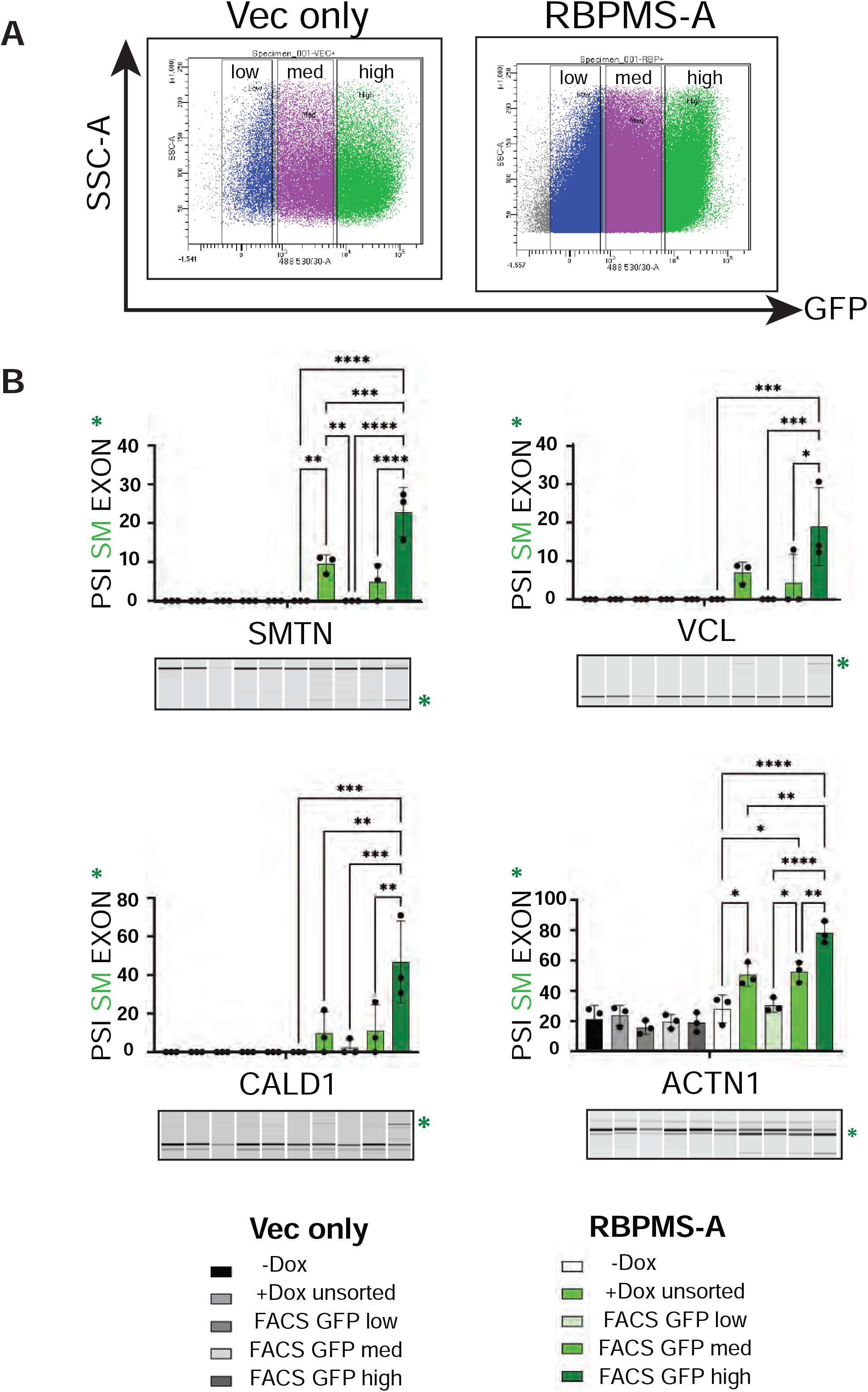
FACS sorted neural crest–derived VSMC fractions for GFP intensity show differential splicing patterns in a RBPMS dose responsive manner. **A**. Doxycycline treated, neural crest derived Vec and RBPMS-A VSMCs were sorted for GFP intensity using flow cytometry, the indicated fractions were collected and RNA was isolated. **B**. RT-PCR of selected SM-AS targets from the Untreated and Doxycycline-treated unsorted, low, medium and high GFP VSMC fractions of the Vec and the RBPMS-A cells. All the targets tested showed RBPMS dose responsive induction of the SM exon. “Percent Spliced In” (PSI) of the SM exon is graphed (as in Fig 1) and the green asterisk highlights the band with the SM exon in the electrophoresis image. Note that in SMTN, the SM isoform is an exon skipping event, whereas VCL, CALD1 and ACTN1 involve inclusion of an SM exon. 3 independent differentiations of 1 Vec and 1 RBPMS clonal line (1 neural crest lineage at different passages were differentiated independently to VSMCs) were FACS sorted and RT-PCRs performed. Ordinary one-way ANOVA with Sidak correction for multiple comparisons was performed across samples within each cell type i.e. Vec vs RBPMS comparisons were not made. Pvalues <0.0001 ****, <0.001 ***, <0.01 **, < 0.05 *. Only statistically significant comparisons are indicated.

We next performed bulk pre-mRNA sequencing to compare the transcriptomes of untreated (No dox), and doxycycline treated low GFP and high GFP populations from both the RBPMS and the Vec hES-VSMCs (Fig 3A). hES-VSMCs of the NC lineage were used for all subsequent experiments. Differential expression analyses showed that the biggest number of differences (>2000 genes) arose when comparing Vec hES-VSMCs to RBPMS hES-VSMCs suggestive of high background variability between these lines and rendering such comparisons noisy. Hence, we restricted our comparisons (differential expression as well as splicing analyses) internally within control Vector lines or RBPMS lines. Another concern was the level of gene expression changes that might be induced simply by Doxycycline treatment. However, we found that in the Vector hES-VSMCs, fewer than 10 genes showed differential expression between untreated and Doxycycline-treated samples whether high or low GFP populations, suggesting minimal influence of drug treatment. Similarly, untreated RBPMS hES-VSMCs when compared with low GFP/RBPMS cells showed significant differential expression in only 5 genes. However, RBPMS/GFP high hES-VSMCs when compared with RBPMS/GFP low VSMCs or untreated cells showed the highest number of differential expression changes (69 and 82 genes respectively) suggesting that RBPMS was the biggest driver of differential expression in these conditions (Fig 3B, Supplemental table 1, see R shiny App (materials and methods)).

**Fig 3.**
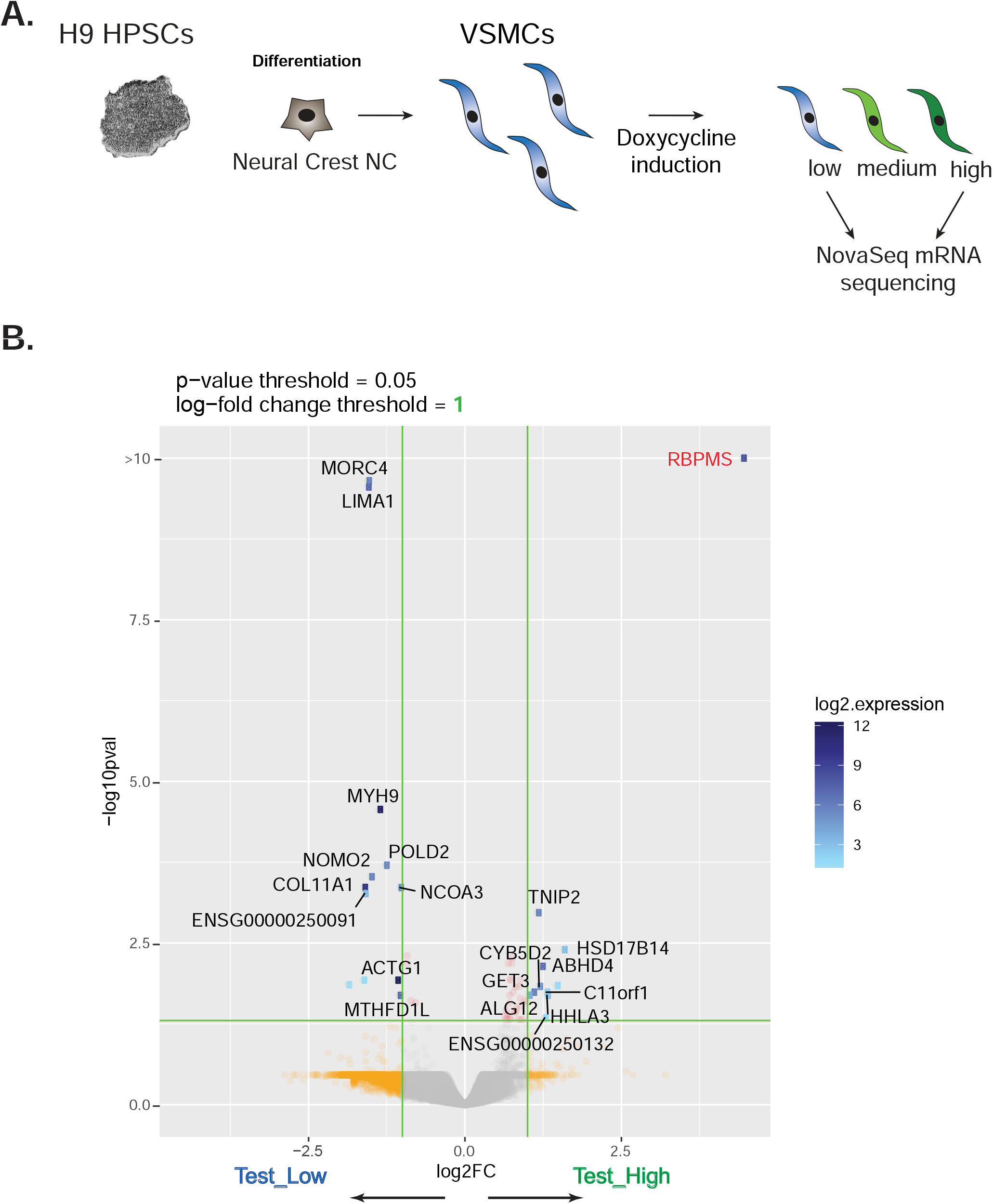
Bulk mRNA sequencing of GFP low and high fractions of Vec and RBPMS-A VSMCs reveals major transcriptomic changes driven by RBPMS. **A**. Schematic of rTTA-ES neural-crest derived VSMCs induced with Doxycycline, sorted for GFP intensity and bulk sequenced using high throughput Nova-seq platform. Both the Vec and the RBPMS-A lines were sequenced. N=4 for the RBPMS-A lines and N=3 for the Vec lines. **B**. Volcano plot showing the differentially expressed genes between RBPMS high and low samples with log-fold change threshold of 1 and p value < 0.05. *RBPMS* is the most upregulated gene.

### RBPMS drives tissue-like contractile SM-AS in hES-VSMCs

We conducted splicing analysis using rMATS with a cut-off of FDR < 0.05 and 15% <u>P</u>ercent <u>S</u>pliced In difference (delPSI). When comparing RBPMS high and low cells, we identified a total of 3673 AS events that were significantly differentially regulated across the 5 categories identified by rMATS - skipped exon, mutually exclusive, retained intron, alternative 3’ splice site and alternative 5’ splice site events (Fig 4A, supplemental table 2). Key events regulated by RBPMS included ACTN1, TPM1 and CALD1 (Fig 4B)– targets that we had previously observed using adult rat PAC1 VSMCs. In each case, RBPMS promoted splicing patterns akin to contractile tissue VSMCs corroborating our findings from the rodent models [6, 7]. Our stringent filters for expression (junction counts) and splicing change - delPSI also meant that certain key targets such as the transcription factor MYOCD (low expression in hES-VSMCs) and meta-VCL (statistically significant but under 15% delPSI) did not pass our cut-offs. However, sashimi plots revealed clear changes in splicing of VCL (Fig 4 associated supplementary A) toward the SM isoforms. In the latter case, although the delPSI value is below the cut-off (∼10% delPSI comparing RBPMS high and low samples), it is important to note the significance of the event because Meta-VCL, a marker of mature VSMCs, is induced from ∼0.2% meta exon inclusion in RBPMS low cells to 10% inclusion in RBPMS-high cells, a magnitude of fold change of at least 50 fold (Fig 4 associated supplementary A). Moreover, meta-VCL induction could be clearly observed by immunoblotting (Fig 4 associated supplementary B) and RT-PCR (Fig 2 and Fig 2 associated supplementary), and even in adult aorta the exon is only included to ∼30% (Fig 4 associated supplementary A). Finally, the number of significant events when filtered by the same criteria were far fewer when comparing 1. GFP high with No-Dox, or with GFP low in the Vec hES-VSMCs and 2. No-Dox and low GFP in RBPMS-hES-VSMCs (Supplemental table 2).

**Fig 4.**
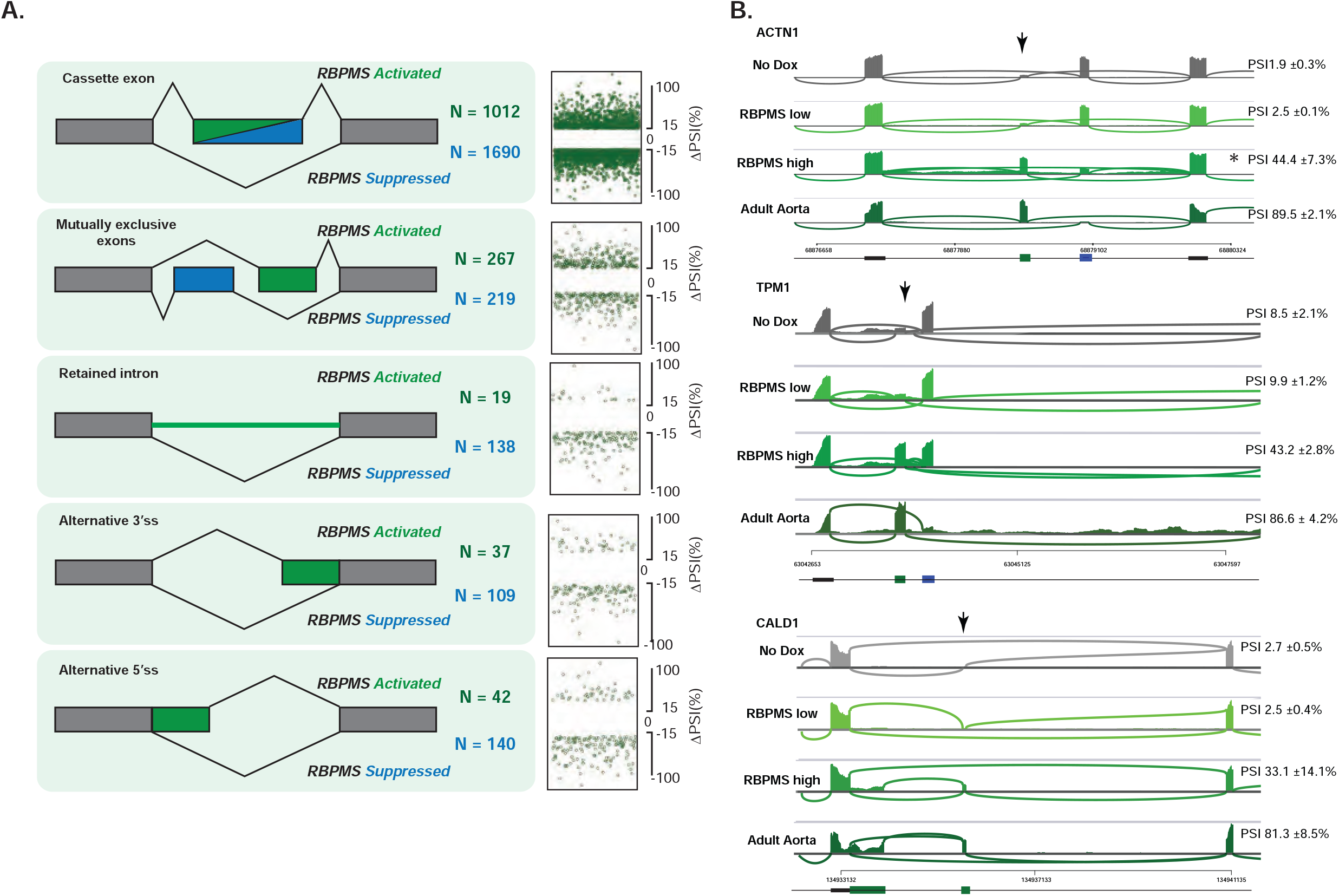
rMATS splicing analysis of RBPMS high and low VSMCs reveals a network of splicing changes induced by RBPMS over-expression. **A**. Splicing patterns altered upon RBPMS over-expression cover >3000 high confidence (FDR < 0.05, |deltaPSI| >= 15%) events across 5 major alternative splicing categories. DeltaPSI represents the change in Percent Spliced In or inclusion of an individual “event” or exon in the final transcript between the RBPMS high and RBPMS low cells. Schematics describe the type of splicing seen in each category whether RBPMS – activated (increased PSI with RBPMS i.e. positive deltaPSI) or RBPMS – repressed (decreased PSI with RBPMS i.e. negative deltaPSI). **B**. Sashimi plots showing expression of the RBPMS-regulated exons and the flanking constitutive regions in *ACTN1, TPM1* and *CALD1*. The SM exon is indicated with an arrow. The No Dox (untreated), low RBPMS, high RBPMS and adult human aorta tissue [54] tracks are shown. A clear increase in the SM exon or isoform is observable in the RBPMS high tracks compared to the controls. PSI values represent the SM exon inclusion percentage +/-standard deviation across N=4 samples for No Dox, RBPMS low and high samples and N=3 for adult aorta samples. In the case of mutually exclusive SM exon in ACTN1, * indicates the discrepancy in PSI values predicted by rMATS and the exon expression visual in the tracks. The trend in increase in SM exon PSI however, remains the same.

We next sought to establish whether the RBPMS-directed alternatively spliced exons showed evidence of regulation by direct binding. To this end, we searched for enrichment of putative RBPMS-motifs using on the differentially spliced cassette exons and 250 nucleotides of their flanking introns. We defined optimal RBPMS motifs as pairs of CAC triplets separated by spacers ranging from 1 to 12 nucleotides reflecting RBPMS binding RNA as a dimer [51]. Indeed, we found a significant enrichment of the RBPMS motif on a subset of transcripts undergoing RBPMS-induced alternative splicing events suggesting that RBPMS directly binds and directs the splicing of its target exons (Fig 5A). Moreover, we detected a positional influence of RBPMS binding on exon splicing. RBPMS motifs were highly enriched in the downstream intron close to the 5’ splice site of exons that were significantly included upon RBPMS over-expression and somewhat less so in the upstream intron. However, exons that were skipped upon RBPMS over-expression showed the presence of RBPMS motifs primarily on the exon itself and in upstream intron overlapping the 3’ splice site (Fig 5A). This pattern is similar to what we have observed previously for RBPMS regulated transcripts in rat models [7] and is also consistent with the position-dependent regulatory rules of many well-known splicing factors [52]. To corroborate this further, we took a more agnostic approach. We performed k-mer (8-mer) enrichment analysis on these differentially spliced targets followed by alignment and hierarchical clustering of the 8-mers that were significantly enriched (p<0.01). This approach yielded consensus 5-mer motifs of highly represented sequences residing in the regulated exons and their flanking introns. In agreement with the specified motif search results, the top enriched 5-mers emerged as CAC-rich (Fig 5 associated supplementary A) particularly in the downstream introns of RBPMS-included exons (Supplemental table 3). Overall, we could deduce RBPMS binding to at least of subset of its target transcripts through motif search alone.

**Fig 5.**
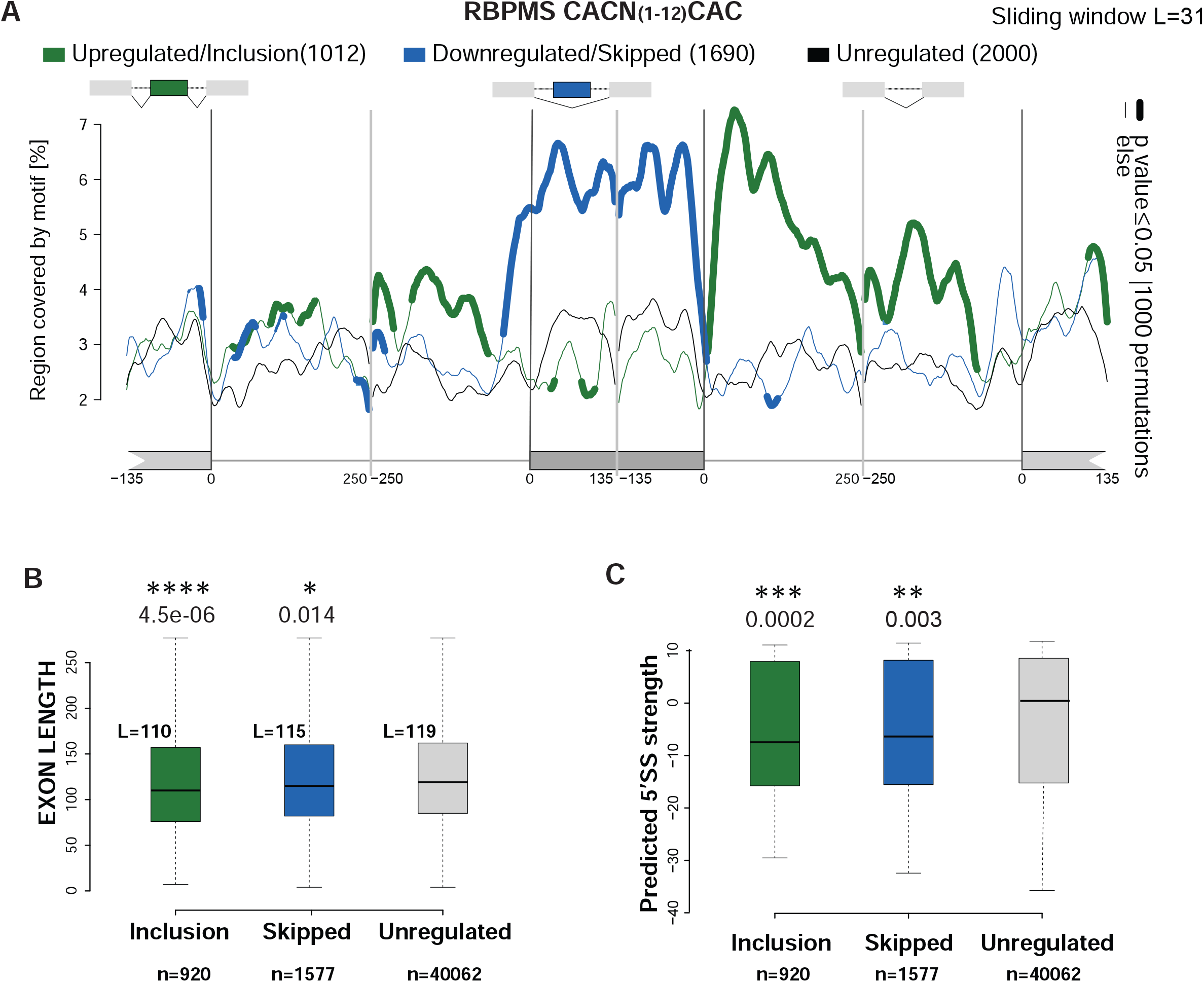
RBPMS regulated exons show enrichment of CAC clusters indicating position-dependent splicing activity. **A**. RBPMS motifs are significantly enriched primarily in the downstream intron of RBPMS-activated exons and on and immediately upstream of RBPMS-repressed exons. RNA maps motif enrichment search with a sliding window length of 31 nucleotides using MATT for the RBPMS binding motif defined as CAC clusters with spacers of length ranging from 1 to 12 nucleotides. The test exons (cassette exon category) were either those activated (increased inclusion – n=1012, green) or repressed (increased skipping – n= 1690, blue) by RBPMS selected using a threshold of 15% or more splicing difference and FDR <0.05. The background exon set were a randomly selected subset of 2000 non-regulated events (FDR > 0.1 and splicing difference < 10%). Statistically significant (permutation test 1000 iterations - p <= 0.05) occurrences on the regulated exon (135bp from the upstream and downstream ends), 250 bp of the flanking introns and 135 bp the flanking constitutive exons are marked by the thick, coloured lines with y axis showing the % region covered by the defined motif. **B**. RBPMS-regulated exons are of consistent and significantly lower length than background exons. Analysis of exon properties by MATT. Significant and biologically relevant categories are presented here. Mann Whitney U test. **C**. RBPMS regulated exons show significantly lowered 5’ splice site strengths compared to the background exons. Analysis was with MATT suite using the MAXENT score - HSA model. Mann Whitney U test.

We also queried the properties of the exons regulated by RBPMS over-expression using the Matt suite [53] (supplemental table 4) and observed that both skipped and included exons were slightly but significantly smaller than control unregulated alternative exons (Fig 5B). All regulated exons had significantly weaker 5’ splice sites (Fig 5C) but not 3’ splice sites as determined by the MAXENT score with HSA model (Fig 5 associated supplementary B). RBPMS controlled transcripts also possessed significantly higher GC content compared to unregulated controls (Fig 5 associated supplementary C). These results suggest that the RBPMS sensitive exons tend to be smaller and their weaker donor splice sites along with the preponderance of RBPMS motifs downstream and proximal to the exon, indicate that they possibly rely on RBPMS to direct the splicing machinery to induce their inclusion.

### RBFOX2 is a potential co-regulator of RBPMS in driving SM-AS

One notable 5-mer sequence that emerged from the k-mer enrichment analysis was GCATG – the optimal motif associated with binding of the RBFOX-family of splicing regulators, which was significantly enriched downstream of RBPMS activated exons (enrichment score = 2.02 pvalue = 0.001). RBFOX2 is the only member of this family that is highly expressed in vascular tissues. Our hES-VSMCs express detectable levels of RBFOX2 (Fig 6 associated supplementary B). We therefore searched for additional clues for the presence of RBFOX2 binding and activity *in silico* in the splicing network altered by RBPMS in hES-VSMCs. We scanned for the occurrence of the RBFOX2 motif on the RBPMS-sensitive exons and their flanking introns and detected a significant enrichment downstream of RBPMS-activated exons and immediately upstream of the 5’ splice site of RBPMS-repressed exons (Fig 6A). Importantly, this pattern coincided with the incidence of RBPMS associated CAC clusters on these regions (Fig. 5A). To address whether the RBPMS and RBFOX2 binding sites are enriched on the same transcripts, we looked for co-occurrence of RBPMS and RBFOX motifs. Using this approach, we could indeed detect significant enrichment of both RBFOX2 and RBPMS motifs occurring as close as 25 nucleotides of each other on the same transcripts – enriched on RBPMS-repressed exons and in the downstream introns of RBPMS-activated exons in hES-VSMCs (Fig 6B). The specificity of association between the two motifs was demonstrated by the minimal enrichment when a single nucleotide change was made to the RBFOX motif (Fig 6. Associated supplementary A). In addition, we performed similar searches for RBFOX2 and linked RBPMS-RBFOX2 motifs and confirmed their enrichment on RBPMS-sensitive exons and introns that we had previously identified in the rat PAC1 VSMC model system under both RBPMS over-expression and knock-down conditions (data not shown) [7]. These results suggest that RBFOX2 and RBPMS might functionally cooperate in exon splicing regulation in VSMCs. To test this possibility, we knocked down RBFOX2 either in combination with RBPMS over-expression or in its absence in RBPMS hES-VSMCs and Vec hES-VSMCs, and analysed AS by RT-PCR. In the SM-AS exons tested, RBFOX2 depletion was able to at least partially reverse the effects of RBPMS over-expression resulting in the loss of contractile splicing patterns (Fig 6C). In the absence of RBPMS (Dox negative hES-VSMCs), RBFOX2 depletion again resulted in the loss of basal hES-VSMC patterns (Fig 6C). This suggests that RBFOX2 is necessary but not sufficient for contractile SM-AS regulation. Rather, it is possible that RBFOX2 is an important cofactor for RBPMS requiring a functional, perhaps physical interaction with RBPMS to promote SM-AS in healthy arterial VSMCs.

**Fig 6.**
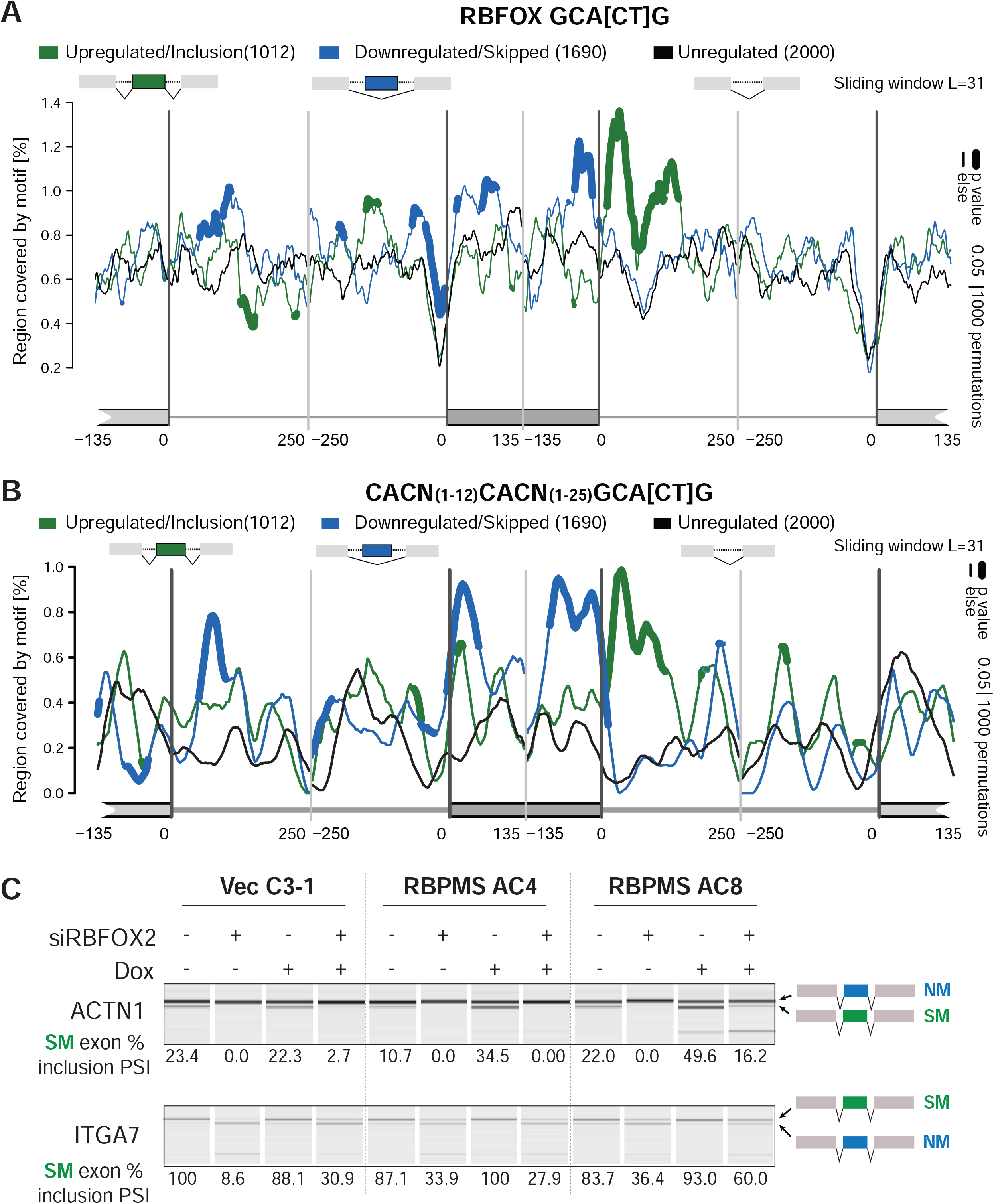
RBFOX2 is a co-regulator of RBPMS at multiple alternatively spliced events in the SM-AS network. **A**. RNA maps of RBFOX motif (GCAC/TG) enrichment around RBPMS regulated and control exons. Analysis as in Figure 5A. **B**. RNA maps showing co-occurring RBPMS and RBFOX2 binding motifs on RBPMS-regulated exons and their flanking introns. **C**. RBFOX2 knock-down experiment with either targeted or control non-specific siRNA treatment of hES-VSMCs cultured with or without 0.2ug/ml Doxycycline. Data are shown for hES-VSMCs derived via neural crest lineage from 1 vector control (Vec) hESC line and 2 RBPMS hESC clones. Each differentiation was performed once. Electropherograms show RT-PCRs of selected targets *ACTN1* and *ITGA7* for the SM exons/isoforms. Quantitated Percent Spliced In (PSI) shown below. RBFOX2 depletion largely decreased endogenous SM splicing in *ITGA7* and in *ACTN1* and substantially reverted the RBPMS-induction of SM exon in *ACTN1*.

### RBPMS-directed SM-AS affects functionally critical gene groups for VSMCs

To investigate the potential biological consequences of RBPMS regulated AS we performed gene ontology (GO) analyses of affected genes. We used a delPSI cut-off of 30% (rather than 15%) to obtain a more stringent network with high confidence AS events for this analysis. Multiple platforms of analyses including GOprofiler (Bioconductor R – Fig 7A) and protein-protein interaction analysis via StringDB (Fig 7B, Fig 7 associated supplementary) yielded terms such as actin binding, focal adhesions and contractile actin filaments as being highly enriched in RBPMS splicing targets. Importantly, GO analyses of the differentially expressed genes induced by RBPMS over-expression in hES-VSMCs did not feature these or any other significant biologically relevant terms (data not shown). These results suggest that RBPMS is capable of remodelling the actin cytoskeleton, potentially to suit the contractile state of VSMCs via changes in splicing patterns rather than by modulating the mRNA abundance of the various component genes. To put the RBPMS regulated splicing network in the context of the AS network of adult human aortic VSMCs, we mined bulk mRNA sequencing datasets of normal adult human aorta [54]. We performed rMATS analysis comparing RBPMS-low hES-VSMCs with the adult aorta and obtained 1737 significantly alternatively spliced skipped exon (SE) events with at least 15% delPSI. Encouragingly, this network showed enrichment of terms including actin cytoskeleton and focal adhesion, similar to the RBPMS regulated AS network (Fig 8 associated supplementary A-C). To delineate the influence of RBPMS in this network, we isolated the commonly differentially spliced events between a) the RBPMS-induced splicing programme in hES-VSMCs (RBPMS high vs low comparison), and b) the Adult aorta vs RBPMS low hES-VSMCs comparison. This yielded 234 SE events that were significantly differentially spliced (FDR <0.05 and |delPSI| >= 15%) across both data sets (Fig 8A). Strikingly, clustering of these events by PSI (percent spliced in) (Z-score normalised shown in heatmap) values showed that RBPMS induced the mature contractile splicing pattern in 75% (175 of 234) of the events (Fig 8B, Supplemental table 5). Of these events, 133 were activated by RBPMS (Group 2) and 42 were repressed (Group 1). In both cases RBPMS overexpression in hES-VSMCs brought the PSI of regulated AS events close to the value found in tissue VSMCs (Fig 8B). The two smaller groups of events with discordance between RBPMS regulation and tissue *vs* hES-VSMC regulation might represent events that are not authentic contractile VSMC markers but instead arise from the comparison of tissue with cultured cells (Fig 8 associated supplementary D). These genes were also not associated with any specific GO terms and a minimal protein-protein interaction network (Supplemental table 5). In contrast, GO and protein network analyses of the 175 events where RBPMS promoted tissue like VSMC patterns showed an enrichment of terms pertaining to actin cytoskeleton organisation and focal adhesions – all connected to smooth muscle contractility and similar to the GO terms enriched in the SM-AS network (Fig 8 C-F). Interestingly, of the 144 genes containing the 175 RBPMS-regulated SM-AS events, 21 (Fig 8C, Supplemental table 5) were associated with super-enhancer regions defined in human aorta tissue (703 total in aorta -dbSUPER database) [55-57], a significant overlap (p = 9.7e-09, pHyper R). This indicates that RBPMS regulates AS events in genes that are crucial for mature VSMC identity.

**Fig 7.**
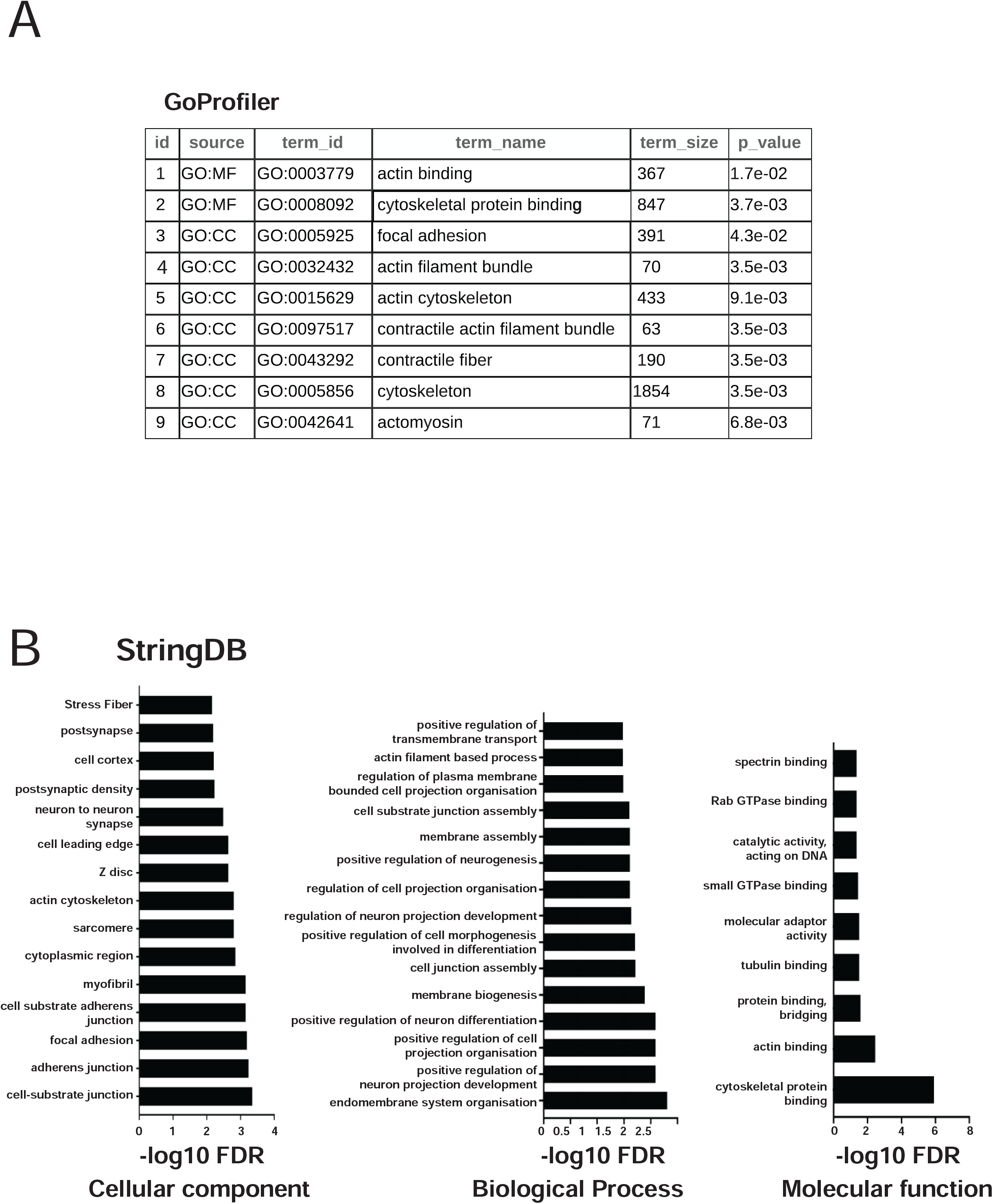
Gene Ontology and protein-protein network analyses of the RBPMS –regulated cassette exons shows functionally critical gene networks for VSMCs. **A**. GOprofiler analyses of the RBPMS-regulated genes (RStudio) using custom background gene sets (all expressed genes in hES-VSMCs). Enriched terms include actin cytoskeleton and focal adhesion components across the Cellular Component (CC), Molecular Function (MF) and Biological Process (BP) categories. Regulated exons were defined as those showing >= 30% splicing differences and FDR <0.05 (Benjamini-Hochberg) **B**. Protein-protein interaction network analysis with StringDB (string-db.org) (whole genome background). Top 15 GO terms from the categories – Cellular Component, Molecular Function and Biological Process are shown, arranged by –log10 FDR values (high to low) are shown. These were filtered limiting to those terms that contained at least 25 genes in the background set in that category and had a strength (log 10 observed/expected) of 0.3 or more.

**Fig 8.**
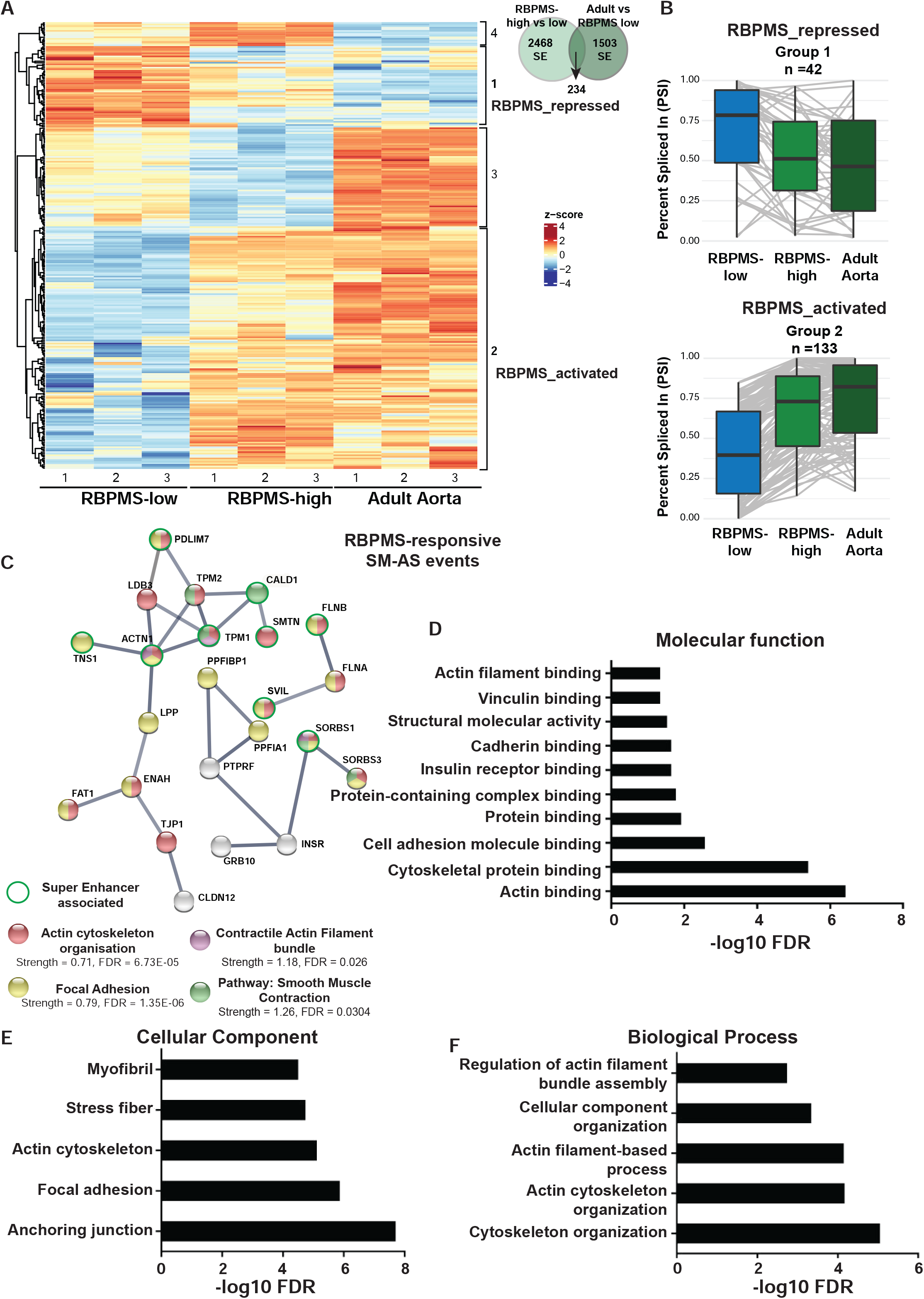
RBPMS promotes tissue VSMC AS patterns. **A**. Heatmap showing events of the cassette exon category significantly (delPSI >=15% and FDR <0.05) alternatively spliced between human aorta tissue and hES-VSMCs that are also regulated by RBPMS. 4 major clusters are identified based on relative splicing patterns between the adult tissue, RBPMS-low and RBPMS-high hES-VSMCs. Clusters 1 and 2 identified as RBPMS-repressed or activated respectively include events where RBPMS regulation matches aorta tissue splicing. Venn diagram shows a total of 234 events detected by rMATS as commonly alternatively spliced between the RBPMS high vs low hES-VSMCs comparison and the adult tissue vs RBPMS-low hES-VSMCs. **B**. Box plots summarizing the relative splicing pattern of the events included in clusters 1 and 2 of the heatmap in A. **C**. Protein-protein interaction network with StringDB of the RBPMS-regulated aorta SM-AS events from B combining genes from clusters 1 and 2. High confidence nodes of interaction scores 0.7 or more (whole genome background) are shown covering key factors involved in the contractile, focal adhesion and actin cytoskeletal machinery. Key shows color code for the nodes and the enrichment strength and FDR values (Benjamini-Hochberg). Nodes outlined with the green circles indicate genes associated with super-enhancers in the human aorta tissue (https://asntech.org/dbsuper/). **D-F**. Bar plots showing top enhanced enriched terms of clusters 1 and 2 in the Molecular Function, Cellular Component and Biological Process categories respectively. The terms enriched are represented with –log10 FDR and filtered from StringDB analyses for term strength (log 10 observed/expected) of 0.1 or more and FDR <= 0.001.

### RBPMS over-expression alters proliferation and motility in hES-VSMCs

Given that focal adhesions and cytoskeleton related terms featured prominently in our GO analyses, we sought to examine phenotypic features of RBPMS over-expressing hES-VSMCs that might be affected by alternative splicing in these complexes. When we compared the motility of RBPMS-high vs RBPMS-low hES-VSMCs in the same Doxycycline-treated population, we observed that RBPMS expressing cells were significantly less motile (lower mean Euclidean distance) than their low RBPMS counterparts when we followed their motion over the course of 8 hours using live cell imaging. We observed this effect in 3 independent RBPMS clonal lines (Fig 9A,B). We confirmed that this effect was not due to GFP expression by comparing GFP high and low cells in Vec hES-VSMCs, which did not differ significantly in their motility (Fig 9 associated supplementary). These results suggest that RBPMS-induced splicing changes are capable of driving phenotypic alterations in hES-VSMCs similar to mature, healthy arterial VSMCs which are also non motile [58]. However, RBPMS over-expression did not cause any perceivable or overt cytoskeletal rearrangements (data not shown) unlike our previous knockdown models in adult rat PAC1 VSMCs [7].

**Fig 9.**
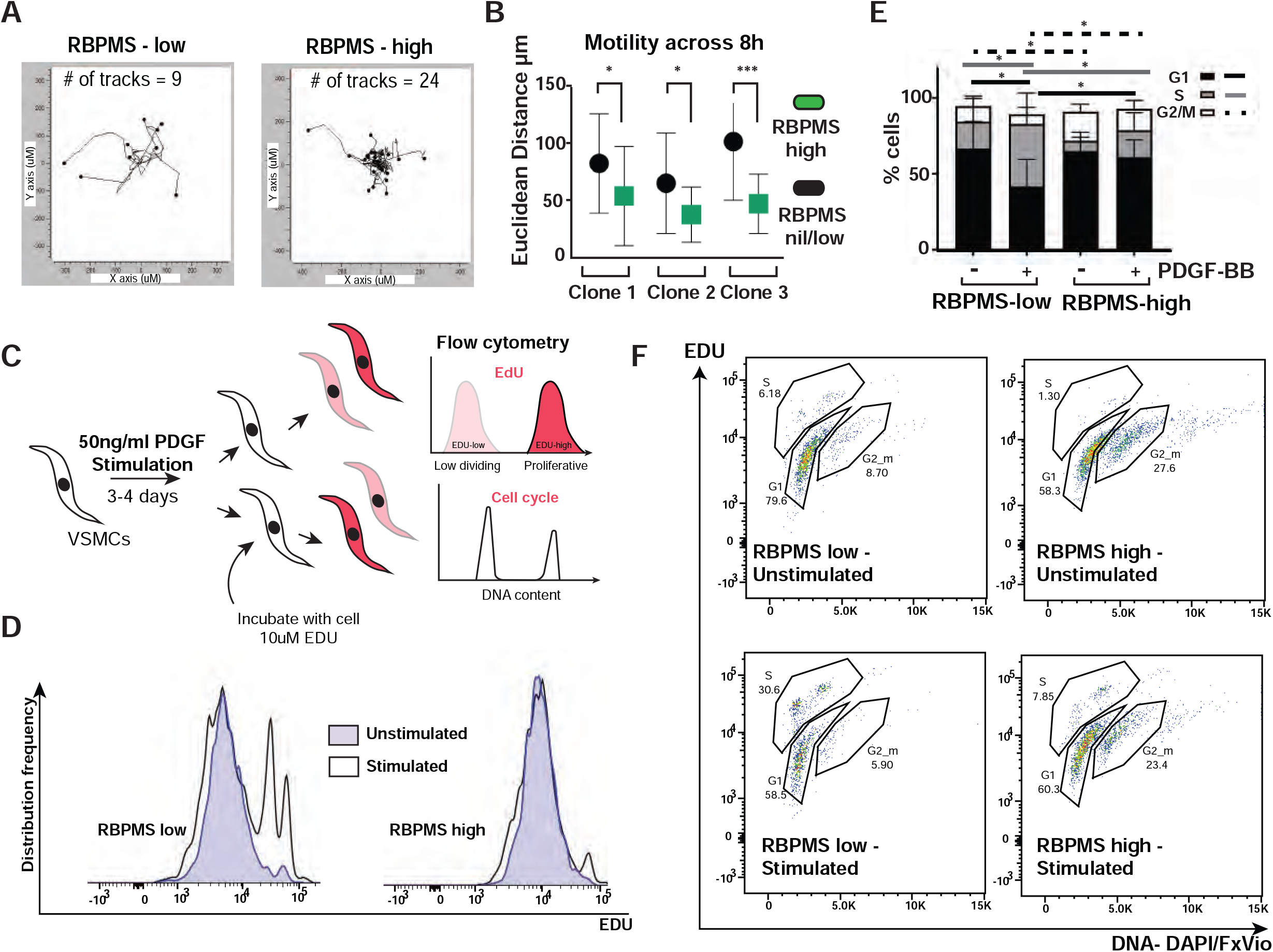
RBPMS overexpression alters hES-VSMC phenotypic behaviour. **A**. Motility of Doxycycline treated RBPMS hES-VSMCs was examined with live cell imaging over the course of 8 hours. Cells were tracked per well with imageJ cell tracker plugin – Manual Tracking. Cell trajectory data were analysed and plotted with the Ibidi Chemotaxis and Migration tool. Representative plots show 9 RBPMS low cells and 20 RBPMS high cells displaying differential motion. **B**. Euclidean distance traversed by 40 GFP positive and 40 GFP negative hES-VSMCs (across 2 wells) derived from 3 independent RBPMS hESC clones is summarised. RBPMS high cells show significantly lower motility in all 3 lines compared to RBPMS low cells (2-way ANOVA multiple comparison without correction independent Fishers LSD-tests p values – clone 1=0.0130, clone 2= 0.0168, clone 3 < 0.0001). **C**. Experimental scheme for hES-VSMC proliferation assay. Doxycycline treated cells were stimulated with 50ng-ml PDGF-B for up to 4 days, labelled with 10uM EDU overnight and assayed for EDU incorporation and DNA content. **D**. Representative distribution frequency histograms (FlowJo population comparison) showing EDU incorporation in PDGF-B stimulated and unstimulated hES-VSMCs gated for low (RBPMS low) and high GFP (RBPMS high). PDGF stimulates EDU labelling in RBPMS low cells but this is substantially reduced in RBPMS high cells. **E**. Summary of the distribution of RBPMS high and low cells (% parent population) with or with PDGF-BB stimulation across the cell cycle. RBPMS high cells do not enter S phase as much as the RBPMS low cells and show higher baseline G2/M populations. Statistical significance of the most meaningful comparisons across the different groups is indicated with black lines (G1 phase differences), grey lines (S phase differences) and dashed black lines (G2/M differences). Fisher’s LSD test without correction for multiple comparisons in 2-way ANOVA. P values are as follows – G1: RBPMS low Unstimulated vs Stimulated =0.0219, RBPMS low vs high Stimulated = 0.0359, S: RBPMS low Unstimulated vs Stimulated =0.0334, RBPMS low vs high Stimulated = 0.0267, G2/M: RBPMS low vs high Unstimulated =0.0145, RBPMS low vs high Stimulated = 0.0140. Data summarise the analyses from 7 independent EDU incorporation assays using hES-VSMCs derived from 2 independent RBPMS hESC clones that could be reliably assayed. Biological replicates within this represented hES-VSMC derivations from at least 2 independent neural crest differentiations per clone. **F**. Gating strategy of a representative hES-VSMC PDGF-induced proliferation assay. These data correspond to the populations compared in the histogram in D. Double staining for incorporated EDU and DNA content with Dapi or Fx-Violet (Invitrogen) allow gating for the populations in different cell cycle stages.

In addition to changes in cytoskeleton related genes, RBPMS also influenced the splicing of a small cluster of cell-cycle associated genes (Supplemental table 2). We queried the proliferative capacity of RBPMS-high cells in comparison with RBPMS-low cells using EDU labelling (Fig 9C-F). We first induced proliferation in the hES-VSMCs by treating them with mitogen PDGF for at least 72 hours. Following this, we labelled the cells with EDU and performed flow cytometry to examine DNA content in addition to EDU incorporation (Fig 9C). We observed that RBPMS-high cells in general displayed lower EDU incorporation when stimulated with PDGF when compared to the RBPMS-low cells (Fig 9D). In addition, a smaller percentage of RBPMS-high cells entered S phase after PDGF treatment (Fig 9 E,F). There was also a higher proportion of RBPMS-high cells in G2/M phase both at baseline and PDGF-stimulated conditions. This result is reproducible in multiple RBPMS-clonal lines. This indicates that RBPMS over-expression alters the proliferative properties of hES-VSMCs pushing them toward a more mature, less proliferative phenotype similar again to the properties of healthy arterial VSMCs.

## Discussion and conclusions

### hES-VSMCs – a model to dissect VSMC maturation and the relationship with SM-AS

The role of a VSMC splicing regulator RBPMS is explored here using *in vitro* human embryonic stem cell derived VSMCs. hES-VSMCs, however, present as inherently immature or potentially foetal-like in their molecular profiles. For example, the differentiation from embryonic stem cells to VSMCs stimulates increase in levels of identity markers such as *CNN1, TAGLN* and *α-SMA* but advanced markers of VSMC maturity such as MYH11 (*SM-MHC*) tend to remain below detection levels [48]. Similarly, hES-VSMCs express basal levels of SM-AS isoforms in several transcripts (e.g *ACTN1*), however, these are variable and do not match those of tissue VSMCs (Figs 1 and 4). Nevertheless, certain genes like *ITGA7* almost exclusively present the SM isoform in hES-VSMCs having switched early on during the differentiation protocol (Fig 1, 6 and data not shown). These results suggest that a push toward SM-AS is incurred during differentiation to hES-VSMCs, however the complete transition to and the maintenance of the SM-AS isoform ratios observed in mature, adult tissue VSMCs, require additional potent, possibly switch-like regulators. We hypothesized that RBPMS is one such critical switch that effects the transition into a mature contractile state via SM-AS induction. Given that the hES-VSMCs do not express RBPMS endogenously, this set the stage to investigate its role in hES-VSMC maturation via over-expression.

### RBPMS is a potent driver of contractile SM-AS akin to mature VSMCs

With an internally controlled, Doxycycline inducible system we show that, RBPMS drives splicing patterns similar to mature tissue-like VSMCs in multiple targets accounting for a critical subset (∼10%-Hypergeometric testing of the overlap showed statistical significance p = 2.0e-35– pHyper R) of the splicing differences seen between adult aorta tissue and hES-VSMCs. This included dramatic increases in the expression of heavy-CALD1 and meta-VCL (Fig 4 and supplementary), acto-myosin crosslinkers both known to be associated with mature, arterial tissue VSMCs as markers of contractility [8-10]. Because this subset of the RBPMS-driven hES-VSMC splicing network comprised functionally relevant, aorta-tissue super-enhancer associated genes (Fig 8) that are involved in various aspects of focal adhesion, actin cytoskeleton structure and contractile function, we are confident that it represents a proportion of the *bona fide* human SM-AS programme (adult human aortic tissue i.e. mature VSMC spliceome) that is controlled by RBPMS.

There are, however caveats to this approach. First, the human SM-AS programme cannot be completely inferred by directly comparing human tissue VSMC splicing profiles to hES-VSMCs or any extant cell culture VSMC model. This is because artefactual molecular differences arising from culture conditions, matrix stiffness and other non-physiological factors will obscure differences reflective of the true VSMC phenotypic state or of the effects of specific genetic perturbations. Second, to properly map the mature/differentiated adult SM-AS network, we would need to compare its splicing profile with a cognate phenotypically switched network, ideally *in vivo* during, for example, injury response [3, 59]. This is achievable in rodent systems but very challenging to model with human VSMCs. Third, the hES-VSMC model explores the splicing activity of a single isoform – RBPMS-A, overexpressed at non-physiological levels (Fig 4 associated supplementary C). RBPMS, (by virtue of its own splicing), is physiologically expressed as multiple transcript isoforms with at least 2 main (A and B) protein isoforms that we have observed to possess differential activity [7, 60]. To compound this, RBPMS activity is also regulated by post-translational modifications [61]. It is possible that in mature tissue VSMCs, the complexity conferred by these additional splice forms and also post-translational modifications [61] fine-tune the regulation of SM-AS by RBPMS. Also, it is possible that a large proportion of the >3000 ASEs that we observe in hES-VSMCs under high RBPMS doses, represents events not directly related to SM-AS but regulated through artificially high RBPMS-A activity. However, by focusing on the overlap of splicing events that are regulated congruently by RBPMS overexpression and in adult aorta, we were able to circumvent the limitations of the individual comparisons and identified a core set of RBPMS regulated AS events that regulate a tight network of proteins involved in cell motility/contraction and cell adhesion (Figure 8). To fully delineate the role of RBPMS in mature VSMC phenotype, systematic, tissue-specific RBPMS-knockout mouse models are required.

The influence of RBPMS on transcriptional changes is minimal in our model. In the case of certain targets such as *PTBP2*, the increase in mRNA abundance is directly linked to splicing of an NMD exon [62] triggered by RBPMS (Supplemental table 2). What we do not observe upon RBPMS over-expression, are significant differences in levels of mature contractile VSMC transcriptional markers (Supplemental table 1, Shiny App) such as *SMTN–B* and the Myocardin (*MYOCD*) and Serum Response Factor (SRF) dependent CArG box genes - *CNN1, α-SMA, SM-MHC* (MYH11) [63, 64]. Indeed, in hES-VSMCs, the effect of RBPMS seems largely confined to splicing regulation of expressed genes. A critical link between RBPMS-regulated SM-AS and the mature VSMC transcriptional network is MYOCD. A recent study indicated that MYOCD transcriptionally upregulates *RBPMS* in VSMCs leading to SM-AS induction upon over-expression [65]. We have shown that RBPMS, in turn, activates splicing of the VSMC-specific exon 2a in *MYOCD* [7], which allows for the generation of an N-terminal truncated protein that has been shown, *in vitro*, to act as the optimal co-factor with SRF to drive expression of the CArG box genes when compared to the non-VSMC isoform (exon 2a skipped) [19]. It is therefore likely that there is a feedback mechanism operating between splicing and transcriptional regulation in mature VSMCs due to the connection between MYOCD and RBPMS, although, the degree of cross talk between the overall pathways that control the transcriptional and post-transcriptional components of the mature VSMC transcriptome remain unknown.

### Mechanisms of RBPMS-driven splicing - RBFOX2, an essential co-factor in a subset of RBPMS-driven SM-AS

We found a significant enrichment of CAC-motifs on and around the alternatively spliced exons in a significant subset of the RBPMS-regulated events indicating the potential for splicing regulation by direct RBPMS binding at these sites along with position-dependent splicing activity – a property typical of several splicing regulatory factors. A possibility is that RBPMS uses co-factors to regulate the splicing of various targets. Our detailed motif analyses on RBPMS-regulated alternatively spliced exons and their flanking introns flagged the GCAC/TG motif, showing coinciding or “linked” occurrence with RBPMS binding sites (Fig 6 and associated supplementary). This motif is highly specific for the RBFOX family of RNA binding proteins - 1, 2 and 3, that amongst a myriad of other functions, are best known as highly versatile splicing regulators [24, 66]. While RBFOX3 is primarily restricted to neuronal tissues, both RBFOX1 and RBFOX2 have been shown to be highly expressed and active in skeletal and cardiac muscle during development and their deregulation has been implicated in the etiology of several cardiac pathologies [24]. However, in smooth muscle rich tissues such as the aorta and bladder, where RBFOX2 is predominantly expressed (GTEX), the role of the RBFOX family is not well understood. We only have limited knowledge of the splicing activity of RBFOX2 in vasculature from its role in the endothelium in mediating the response to low blood flow [38] and its regulation of the splicing of the Calcium channel CaV1.2 in VSMCs implicated in the pathology of hypertension alongside RBFOX1 [39, 67].

The co-occurrence of RBFOX2 motifs with RBPMS, suggested a co-regulatory function in SM-AS. Our RBFOX2 depletion experiments confirmed this and revealed it to be a necessary co-factor for SM-AS induction in hES-VSMCs in a panel of RBPMS-targets tested, although the magnitude of its impact varies between different transcripts and the effects were quite variable across hES-VSMC clones. For example, loss of RBFOX2 largely reversed SM-AS, either RBPMS-induced in *ACTN1* or endogenous as in *ITGA7*. We do not think RBFOX2 itself fits the criteria to be classed as a master regulatory splicing factor in VSMCs primarily because of its wide expression and activity in other tissues and because, unlike RBPMS, its expression is stable and not dynamically regulated during phenotype switching (data from [7]). Instead, it seems to act as an essential (but not necessarily sufficient) cofactor with RBPMS functioning via defined VSMC-specific interactomes to specify SM-AS patterns. RBPMS is known to interact with RBFOX2 (Biogrid) in high-throughput experiments, and we have recently found that the two proteins interact directly in cell-free splicing assays (Yang et al, manuscript in preparation). It is not fully clear yet how RBFOX2 and RBPMS physically interact in hES-VSMCs to contribute to SM-AS and how this translates to SM-AS in tissue VSMCs. It is clear from our findings that these proteins have an important functional interaction that is at least partly responsible for the mature VSMC spliceome.

Apart from RBFOX2, it is likely also that RBPMS maintains other interacting partners for driving SM-AS via the formation of large complexes of splicing factors similar to the LASR complex [68] that direct the coordinated post-transcriptional processing of numerous pre-mRNAs in VSMCs. It is likely that RBPMS, whose C-terminus encodes an intrinsically disordered region, interacts in such a complex without direct binding and directs other splicing factors (Yang et al, manuscript in preparation). As a master regulator, we expect that RBPMS will alter the splicing of other RNA binding proteins. RBPMS altered the splicing of at least 160 other proteins classified as RNA binding in RBPbase, including splicing associated factors such as MBNL1 and SNRPA1 (Fig 7 associated supplementary, Supplemental table 2). It is therefore likely that the activity of the altered splice isoforms of these RNA binding proteins also contribute to SM-AS signatures by binding their target transcripts. For example, we have previously shown in the rodent model that RBPMS-regulated MBNL1 splicing affects the splicing of its own target, *NCOR2* [7]. Here also, we observe significant regulation of *NCOR2* (Supplemental table 2) exons in our hES-VSMCs upon RBPMS over-expression.

### RBPMS affects VSMC phenotypic behaviour

We hypothesized that the splicing changes induced in the focal adhesion and cytoskeleton and associated proteins have an effect on multiple phenotypic properties of the cells including the cell’s migration. Focal adhesion proteins mediate complex connections between the cytoskeleton with the extra-cellular matrix affecting the cell’s interaction with its environment and it is likely that alternative splice isoforms of these would mediate differential behaviours. For example, the SM-AS isoform meta-VCL is structurally distinct from VCL and has different properties pertaining to F-actin bundling [69] which bears implications for mechano-transduction, focal adhesion properties and cell motility [70]. RBPMS over-expressing cells were indeed less motile reflective of the low migratory properties of mature VSMCs. To what extent the alternatively spliced isoforms of the focal adhesion complexes contributed to this phenotype will have to be explored. In contrast to the effects of RBPMS depletion in partially differentiated PAC1 cells, where the alterations in AS affecting cytoskeletal protein isoforms correlated with changes in actin cytoskeletal structure, we observed no such change in actin organization upon RBPMS induction in hES-VSMCs (data not shown). Although our over-expression model affected many of the same AS events (in the opposite manner to the knockdown model), it is possible that the splicing changes are necessary but insufficient to detect wholesale actin reorganization towards a contractile state.

Mature VSMCs are quiescent and non-proliferative unless triggered by injury or insult to the vessel wall. hES-VSMCs are themselves proliferative, particularly with mitogenic stimulation with PDGF-B. RBPMS over-expression was able to at least partly suppress the proliferation of hES-VSMCs with perceivable effects on S-phase entry and G2/M. This is an important finding because a recent *in vivo* study by Gan *et al* showed the converse - that the loss of RBPMS in VSMCs during development led to increased proliferation of VSMCs [45], consistent with our observations. RBPMS also increased the inclusion of exon 30 of the epithelial-mesenchymal transition (EMT) associated factor *FLNB* (supplemental table 2),whose skipping is a marker of EMT [71], suggesting that in general in hES-VSMCs, RBPMS can promote a more quiescent, non-mesenchymal state. Phenotypically switched VSMCs are more likely to represent mesenchymal states [72] and that RBPMS is functionally interacting with these processes, implies it is a key regulator of VSMC phenotypic status.

Another major hub possibly influenced by RBPMS includes ECM modulation. RBPMS also targeted the ECM factor *FN1* (supplemental table 2) with increased usage of the EDB exon 25 [73] that has been shown, along with the EDA exon to be essential for embryonic cardiovascular development and in processes like angiogenesis [74]. These exons also maintain complex interactions with the TGF-B pathway, further implicating a relationship for RBPMS with EMT processes in hES-VSMCs. Apart from FN1, several collagen genes show altered splicing in the presence of RBPMS. Although the exact implications of the splicing-induced structural or domain changes are not fully clear, we can speculate that the presence of RBPMS alters the synthetic properties of the hES-VSMCs. Taken together, it is apparent that reach of RBPMS-induced splicing alterations extends beyond actin-cytoskeleton and contractile machinery which predominate the VSMC-relevant GO terms with RBPMS affecting the splicing patterns of key genes in cell cycle and EMT regulatory hubs.

*In vitro* cell culture studies have implicated RBPMS as a regulator of mRNA translation in embryonic stem cells required for differentiation into myocardial lineages [42]. Rbpms homozygous knockout models from both the IMPC (Internation Mouse Phenotyping Consortium - http://www.mousephenotype.org/ [75] - Rbpms^em1(IMPC)H^ allele) and more recently, from Gan et al [45] showed that RBPMS is an essential gene with mice showing pre-weaning/neo-natal lethality with 100% penetrance. Interestingly, this study suggested that the lethal phenotype is a result of under-developed myocardium due to the loss of RBPMS having impaired proliferation in cardiomyocytes at critical embryonic stages by mis-splicing of PDLIM5. This is despite the fact that VSMC proliferation was enhanced in the developing vasculature - the opposite effect from cardiomyocytes - and a failure of the closure of the Ductus Arteriosus after birth i.e. Patent Ductus arteriosus [45]. This is possibly due to the failure to transition to a contractile VSMC phenotype, a necessary stage for ductus arteriosus closure immediately after birth [45, 76]. This bears implications for the importance of RBPMS in the transition from a proliferative to a mature or differentiated contractile state in developing and post-natal vasculature. Single cell studies have indicated that RBPMS is itself a marker of contractile VSMCs [2].We see that RBPMS promoted differentiated, contractile-VSMC like behaviours in hES-VSMCs. We correlate these phenotypic alterations with its induction of the SM-AS programme. However, RBPMS is known to wear many hats – it is a transcriptional co-activator with the AP1 [77] and SMAD [78] signalling, a regulator of translation in embryonic stem cells [42] and associated with RNP granules in oocytes [79]. So we postulate that RBPMS, either directly or indirectly regulates multiple pathways feeding into different cellular processes that are necessary for the induction and maintenance of the contractile VSMC state. RBPMS levels and activity are tightly and dynamically regulated in multiple ways – transcriptional control [7, 60], phosphorylation [61] and even by its own splicing [7, 60, 77] (and data not shown). This suggests that RBPMS, although perhaps not the whole picture, is still a critical regulator of VSMC phenotype of the class of a master regulator. However, to establish a direct relationship between RBPMS-regulated pathway and VSMC phenotypes, more nuanced conditional knockout strategies are necessary in combination with intricate single cell sequencing studies.

## Materials and Methods

### Doxycycline inducible system

The H9-rTTA human embryonic stem cells (hESCs) as described in [50, 80, 81] were used to insert a cassette containing either GFP or RBPMS-A (isoform A [7]) cDNA linked to GFP via a T2A linker sequence under the control of Doxycycline-inducible promoters, into the pUC-AAVS genomic safe harbour locus. Briefly, hESCs were dissociated into single cells and electroporated with plasmids containing Zinc finger nucleases along with the pUC-AAVS constructs bearing the relevant cassettes. Positive clones were selected with 1ug/ml puromycin treatment and isolated and expanded as individual clones of either rTTA-Vec or rTTA-RBP-A hESCs. Clones were eventually expanded and maintained on Vitronectin coated plates and E8 media without puromycin (we did not observe any drift and clones were stable in Puromycin over multiple passages). For differentiation to neural crest lineages, clones were “acclimatised” to 0.1% gelatin coated plates for 2 or 3 passages and then differentiated. Clones were validated for pUC-AAVS locus activity with 1ug/ml doxycycline treatment and GFP induction.

### Vascular Smooth Muscle Cell differentiation

VSMCs were differentiated via the neural crest (NC) or the lateral plate mesoderm (LM) lineages were as described previously in [1, 48, 49]. Briefly, hESCs were seeded 24h prior to differentiation on vitronectin (LM only) or 0.1% Gelatin coated plates in “E8” media containing. Following this, the cells were treated in FSB media for 5 days to direct them toward NC lineage or with a combination of FlyB (36h) and FB (84h) for the LM lineage. For NC, cells were further passaged a minimum of 5 times and a maximum of 15 passages in FSB before use for VSMC differentiations. In the case of LM, once the intermediate was obtained, cells were directly used for VSMC production. LM or NC cells were treated for 12 days in PT media to generate VSMCs which were further matured in SMC media. For Doxycycline induction of RBPMS or Vec VSMC lines, cells were cultured in SMC media containing 0.2ug/ml Doxycycline for a minimum of 5 days before phenotyping and molecular analyses. Media recipes are in supplemental table 7.

### SiRNA knockdown

RBPMS and Vec hES-VSMCs were differentiated from neural crest intermediates and treated with or without Doxycycline (0.2ug/ml) for a minimum of 5 days (to induce RBPMS and GFP expression). Following this, cells were treated with either control (C2) or RBFOX2 siRNA (siRBFOX2) with Dharmafect 1 reagent (Dharmacon-Horizon T-2001-02) according to manufacturer’s protocols. Cells were transfected with 100pmol of Ctrl or anti-RBFOX2 siRNA per well of a 6-well plate for 72 hours and then treated a second time with the same dose for another 48h following which, they were harvested for RNA (Zymo – Directzol kit) and protein. C2 ctrl was used as in [7] and siRBFOX2 – Life technologies silencer predesigned siRNA-ID#136600 -GCCUUUUACUACCAUCCCAtt.

### Splicing assays and RNA sequencing

Vec or RBPMS hES-VSMCs derived via the NC lineage were treated and expanded in SMC media with 0.2ug/ml Doxycycline for at least 5 days before harvest. In parallel, untreated controls – no Dox - were maintained and expanded in SMC media. Cells (both treated and untreated) were harvested by trypsinisation and FACS sorted for GFP intensity into “low”, “medium” and “high” gates. Once collected, the cells were resuspended in either RLT (Qiagen RNeasy) buffer for RNA isolation or in protein lysis buffer (laemmli) and snap frozen until use. RNA was isolated using Directzol (Zymo Research) kits. In general, VSMC lines obtained from multiple neural crest differentiations and/or independent hESC clones were considered as biological replicates. For RNA sequencing of Vec-hES-VSMCs, 2 independent rTTA-Vec clonal lines were used with one clone differentiated into 2 independent NC lines (total 3x). Similarly, for RBPMS-hES-VSMCs, 2 independent rTTA-RBPA-hESC clones were differentiated into 2 separate NC lines each (total 4x). A similar approach was used for RT-PCR validation of splicing. To avoid batch effects, all the RNA samples were frozen upon collection and harvested simultaneously.

For RT-PCR splicing analyses, a minimum of 50ng of RNA was used to prepare cDNA (Roche RT) and splice isoform specific PCRs were performed (Sigma Aldrich High fidelity Taq) using appropriate primers (Supplemental table 8) and resolved using Qiaxcel capillary electrophoresis. Relative quantitation of splice isoforms was done calculating the relative band intensity percentage with Qiaxcel software.

### Antibodies and immunoblotting

Immunoblotting was carried out using standard protocols. Protein samples were separated on 12-15% SDS-PAGE gels and transferred onto PVDF membranes and then probed for expression of specific proteins. Antibodies used were anti-RBPMS (Atlas antibodies HPA056999, 1:1000 dil), anti-RBFOX2 (Atlas antibodies HPA006240 – 1:500 dil), Vinculin (Sigma Aldrich v9264, 1:1000 dil), GAPDH (Sigma Aldrich G9545, 1:2000 dil).

### Library preparation

of poly-A tailed RNA was performed with the NEB PolyA kit (E7490) and the NEB Ultra II directional library kit (E7760). 100ng of total RNA was used and initial quality control was done with Agilent RNA tapestation / RNA bioanalyzer reagents, and with the Qubit RNA HS Assay Kit (Q32855). Quality Control of the libraries was done with Qubit dsDNA HS Assay Kit (Q32854) and Agilent DNA 5000 tapestation reagents. Two randomised pools (of 11 and 10 samples) were sequenced on two lanes of a Novaseq 6000 S1 flowcell, PE150 (Cancer Research UK, Cambridge). Library preparation was conducted by the CSCI genomics facility.

### Analysis pipelines

For all analyses, quality checks were performed using FastQC (https://www.bioinformatics.babraham.ac.uk/projects/fastqc). The alignment to the reference genome (*H. sapiens* hg38, genome assembly GRCh38.p13) was done using STAR version 2.5.2a [82]. Pre- and post-alignment quality checks were summarised using MultiQC. Gene expression counts were generated using featureCounts version v1.6.0 (C). Additional quality checks include MA plots, PCA plots and heatmaps representing the Jaccard Similarity Index (JSI) [83].

The noise identification (on summarised counts and transcripts) and subsequent correction were performed using noisyR [84]. Pearson Correlation was used to determine the signal to noise transition; a correlation threshold of 0.25 between expression profiles across transcripts (transcript approach) and sorted gene expression vectors (count matrix approach), respectively was used for identifying the dataset specific signal to noise value. For the latter, the length of the vectors (sliding windows) was set to one tenth of the total number of transcripts. Both approaches yielded similar signal/noise thresholds (subsequent analyses were based on a noise threshold of 34). All transcripts with abundances less than the signal/noise threshold across all samples were excluded from subsequent analyses. The transcripts with at least one entry above the signal/noise threshold were kept for further analyses. All entries lower than the signal/noise threshold were increased to the threshold, to avoid false positive calls on the DE step.

The normalisation of expression levels was performed using quantile normalisation [85] (using the function normalize.quantiles() from the R package preprocessCore (https://github.com/bmbolstad/preprocessCore)). The differential expression (DE) analysis was performed using the standard functions from edgeR pipeline, version 3.28.0 [86]. The DE analysis was performed on the comparison of Test High against Test Low. The thresholds for DE were |log_2_(FC)| >0.5 and adjusted p-val < 0.05 (adjustment done using the Benjamini-Hochberg method). The differentially expressed genes were summarised in a volcano plot. Enrichment analysis was performed using g:profiler (R package gprofiler2, version 0.2.0) [87], against the standard GO terms, and the KEGG [88] and reactome [89] pathway databases. The observed set consisted of the DE genes, the background set comprised all expressed genes, using the denoised count matrix.

All steps of the analysis can be further explored using a bulkAnalyseR Shiny app [90], created for this dataset. This is available upon request.

### Splicing analysis

Alternative splicing analysis in Ctrl and Test samples was performed comparing GFP high, GFP low or No-Doxycycline samples using rMATS [91] (version 4.1.0) with the BAM files filtered as described above. The splicing profiles of adult human aorta tissue were obtained from GSE147026 [54]. The fastq files obtained from this study were processed and filtered using the same pipeline described above. Similarly noise filtered BAM files were used as input for rMATS. Raw rMATS outputs were then filtered for events that showed statistical significance using an FDR threshold of 5% and an inclusion level difference threshold of 15%. Further filtering on events for which there were at least 80 occurrences in each sample set (either inclusions or skipped junction reads) was applied, to exclude potentially spurious, noisy events.

### Motif analyses

Motif and exon-intron feature analyses of RBPMS-responsive alternatively spliced exons was performed using Matt [53] (v1.2.0). For these analyses only the skipped exon category events (SE outputs from rMATS), being by far, the large majority, were used. The motif corresponding to RBPMS binding sites are CAC clusters defined as CACN(1-12)CAC and the motif corresponding to RBFOX was defined as GCAC/TG. Specified motif searches were performed on the RBPMS-regulated exons and 250nt of the flanking introns using the rna_maps or the test_regexp_enrich tool. Test exons (inclusion/upregulated events and skipped/downregulated events) passed a threshold of FDR<0.05 and Inclusion Level Difference of 15% while background exons (Unregulated) were selected as events with FDR >0.1 and Inclusion level difference of <5%. For running rna maps, the background set was downsampled to a random 2000 events.

For an agnostic approach to identify putative splicing regulatory elements, kmer analysis specifically 8-mer analysis was performed using the test_regexp_enrich tool. Enriched 6mers and 8mers were identified for each comparison, separately for the following variants, following the structure of the Matt pipeline: upstream intron / exon / downstream intron, up-regulation / down-regulation, enrichment / depletion, first / internal / end. All possible 3mers, 4mers, and 5mers were then aligned to the identified enriched 6mers or 8mers to identify recurring short patterns with high enrichment scores for each condition. This analysis was performed in R, utilising the msa package (v1.24.0). The results were summarised using sequence logos visualisation, on information content, as well as heatmaps of the consensus sequences found. The colour gradient of each letter is proportional to the total observed enrichment.

### Gene Ontology analyses of alternatively spliced events

GOprofiler was used with a custom background of all expressed genes in hES-VSMCs for gene ontology with FDR filter of 0.05. For protein-protein network analyses, stringDB (https://string-db.org/)[92] was used with a whole genome background gene set. Typically, the terms presented were filtered for enrichment strength (Log10(observed / expected) threshold of 0.1 and false discovery rate of 0.001. In both analyses, the data set used were the high confidence, cassette exon (SE) events passing a delPSI threshold of 30% in RBPMS high versus low comparisons.

### Cell motility

Live cell imaging was performed with the incucyte system (5% CO2, 96% Rh). Briefly, hES-VSMCs were plated (duplicate wells) and imaged every 2h over the course of at least 8h. 3 independent clonal lines for the RBP-A hES-VSMCs were imaged and 1 clonal line for Vec-hES-VSMCs. Cell tracking was then performed using the imageJ manual cell tracking plugin (http://rsb.info.nih.gov/ij/plugins/track/track.html) whose outputs were analysed with the ibidi cell tracker (Chemotaxis and Migration tool Ver 2.0) software. Analyses were performed blinded.

### Flow cytometry and Proliferation assays

Doxycycline treated hES-RBPMS VSMCs were either treated with 50ng/ml of PDGF-BB or left unstimulated over the course of 72h up to 96h. Cells were then dosed with 10uM EDU overnight, trypisinised, fixed with 4% PFA and analysed for EDU incorporation (Click-iT plus EDU kit – Thermofisher Scientific - C10634) and DNA content (Fx-Violet Thermofisher F10347 or DAPI) on a BD-Fortessa flow cytometer with area parameters enabled. Analysis was performed with Flowjo V10.7.1. A minimum of 1000 events was collected in each sample. Note – the cells were not synchronised for these experiments. For RBPMS intra-cellular staining, cells were fixed in 4% PFA, permeabilized using the BD Biosciences Cytofix/Cytoperm kit (554714), and stained with the RBPMS atlas antibody (same as for immunoblotting – 1:250 dilution) overnight and then with Alexafluor 647 conjugated anti-rabbit secondary antibody.

### Statistics

Statistics were performed and most graphical data were generated with GraphPad Prism ver7 and above.

## Supplemental Tables and shinyApp access

will be available upon request only.

## Figure legends

**Fig 1 associated supplementary 1.**
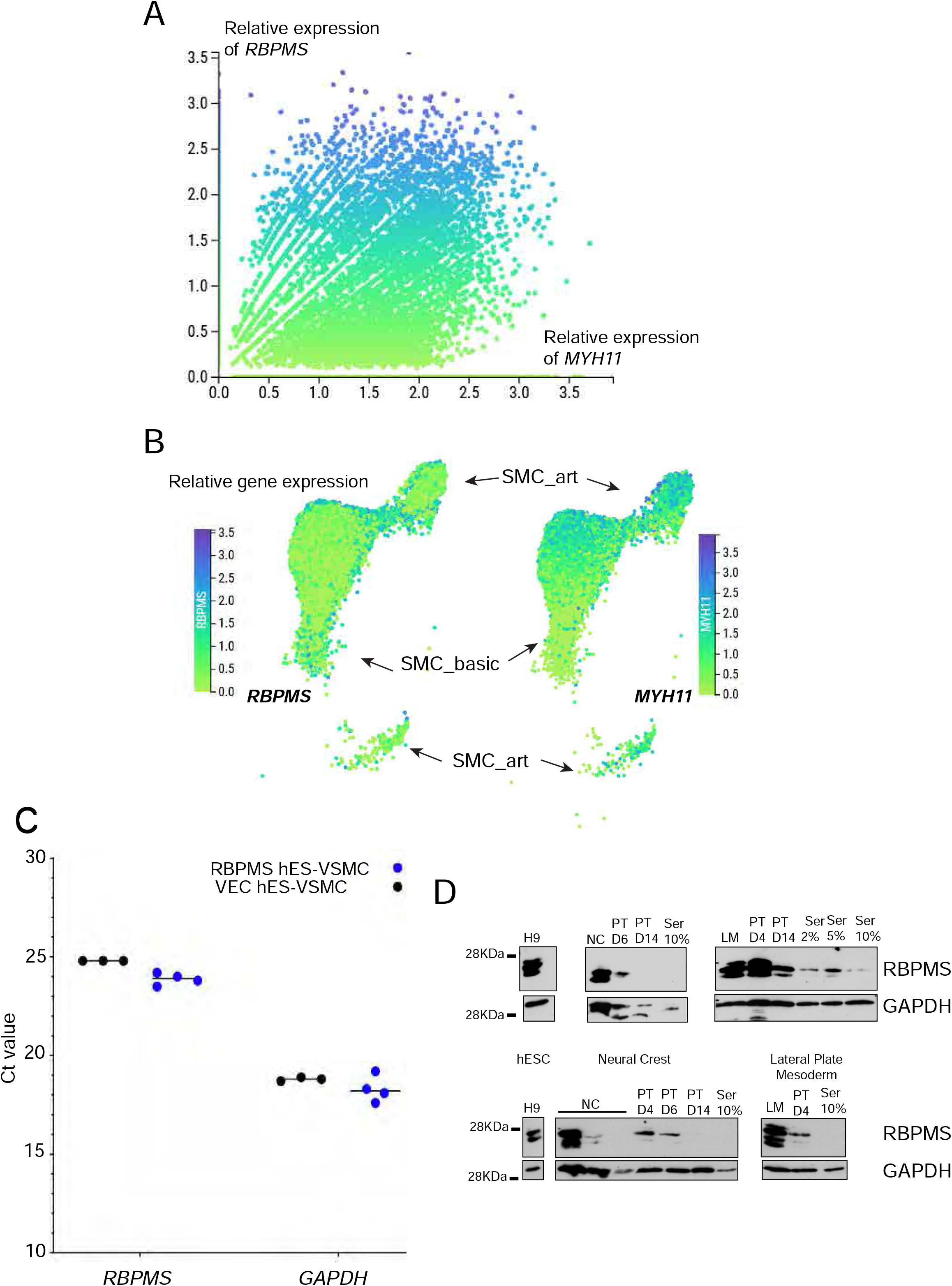
**A**. RBPMS correlates with MYH11 expression in vascular smooth muscle cells. Data are UMAPs of relative gene expression of *RBPMS* and *MYH11* generated using visualization tool CellXgene of the “vascular” data from the human heart cell atlas portal [5]. **B**. UMAPS of the cells classified as smooth muscle show 3 main clusters consisting of SMC-art (arterial) SMC-basic cells that are considered more contractile and proliferative respectively [5]. **C**. *RBPMS* transcripts are expressed at detectable levels in neural crest derived hES-VSMCs. Shown are quantitative RT-PCR data – Ct (amplification cycle) values of *RBPMS* and *GAPDH* in VEC and RBPMS hES-VSMC lines (N=4, derived from 2 independent VEC-hESC and 2 independent RBPMS hES-C clonal lines from multiple neural crest differentiations). These are endogenous *RBPMS* levels without Doxycycline induction. **D**. Immunoblots showing RBPMS levels decrease across differentiation from H9 human embryonic stem cells to VSMCs in 10% serum via the neural crest (NC) and the lateral plate mesoderm (LM) lineages. GAPDH is the loading control. The two panels show two independent experiments.

**Fig 1 associated supplementary 2.**
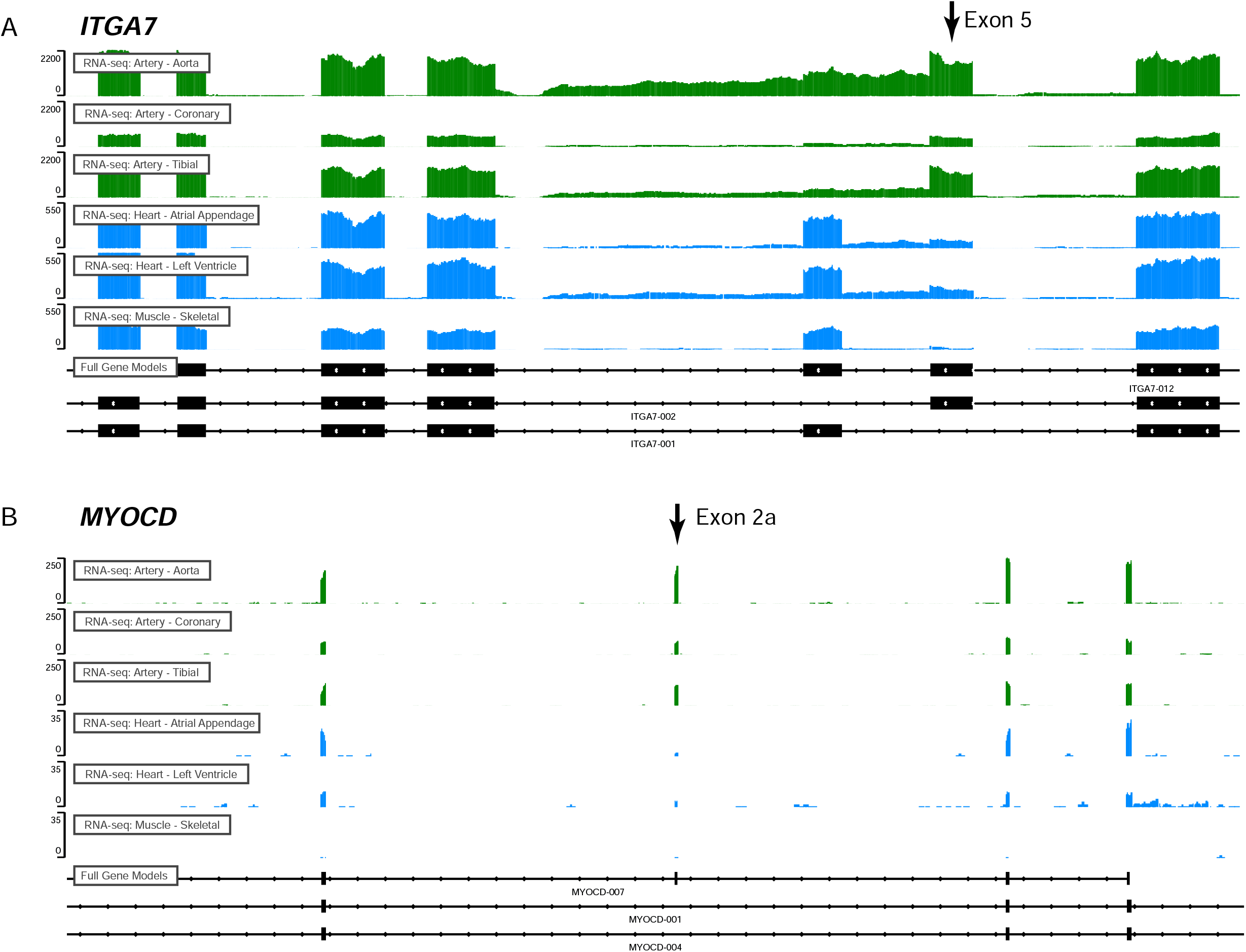
GTEX bulk human adult tissue mRNA-seq IGV tracks showing expression of SM-exons (arrow) in **A**. *ITGA7* and **B**. *MYOCD* in VSMC rich artery tissues – aorta, coronary and tibial (green) and in non-smooth muscle i.e. striated muscle tissues – heart (left and right ventricles) and skeletal muscle (blue). *ITGA7* exon 5 is one of a mutually exclusive pair; exon 6 is used in cardiac and skeletal muscle.

**Fig 2 associated supplementary.**
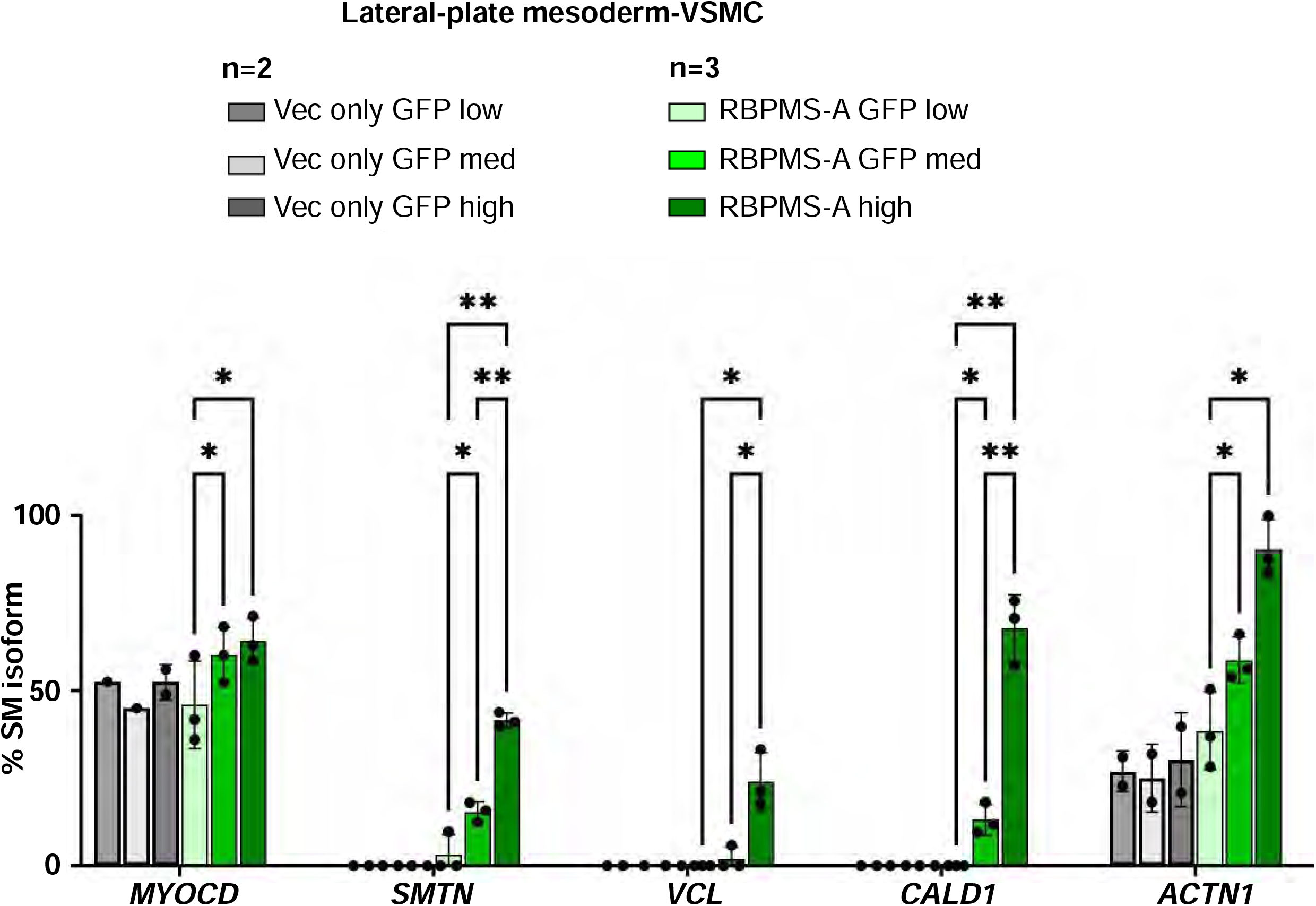
FACS sorted lateral plate mesoderm-derived VSMC fractions for GFP intensity show differential splicing patterns in a RBPMS dose responsive manner. Doxycycline treated, lateral plate mesoderm derived Vec and RBPMS-A VSMCs were sorted for GFP intensity using flow cytometry and the fractions were collected and RNA was isolated. RT-PCR of selected SM-AS targets from the Doxycycline-treated low, medium and high GFP VSMC fractions of the Vec and the RBPMS-A cells. All targets tested showed statistically significant RBPMS dose responsive induction of the SM splicing event. “Percent” of the SM isoform is graphed (as in Fig 1). N=3 for the RBPMS-A line and N=2 for the Vec line where each bioreplicate represents an independent differentiation experiment. RBPMS-A lines were derived from 2 independent clones. Note that in some experiments in the Vec line where GFP intensity is generally higher than the RBPMS-A line, only the high GFP intensity cells could be collected for RNA isolation. Hence statistical analysis was not performed in the VEC samples although the RT-PCR results were reproducible. SM exon inclusion was compared between RBPMS-A VSMCs of low, mid or high GFP gates using 2-WAY ANOVA multiple comparisons without correction (Fisher’s LSD). Pvalues <0.01 **, < 0.05 *. Only statistically significant comparisons are indicated.

**Fig 4 associated supplementary.**
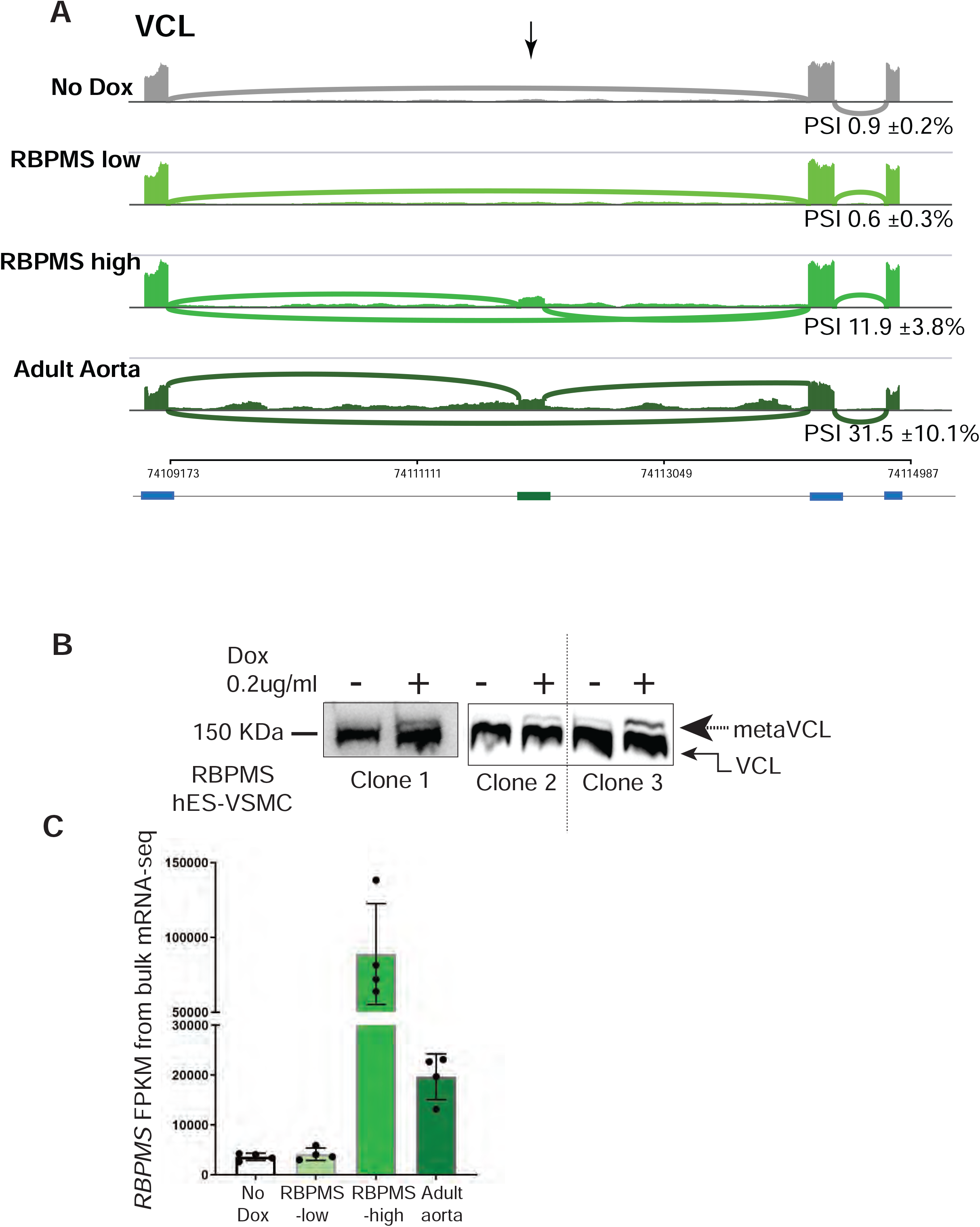
RBPMS induces meta-Vinculin expression. **A**. Sashimi tracks showing meta *VCL* SM isoform (arrow) induction by RBPMS over-expression in a manner similar to adult aorta tissue (obtained from GSE147026) [54]. **B**. Lysates from neural crest derived RBPMS hES-VSMCs either untreated or treated with 0.2ug/ml Doxycycline were separated on a 10% polyacrylamide gel and blotted and probed for Vinculin (VCL) expression. RBPMS over-expression shows induction of the meta-VCL protein isoform matching transcript expression of meta-*VCL* - RT-PCR and bulk mRNA sequencing data. **C**. Graph showing FPKM (fragments per kilobase of exon per million mapped fragments) of *RBPMS* transcripts from bulk mRNA sequencing of RBPMS hES-VSMCs (no Dox, low and high) and adult human aorta tissue [54].

**Fig 5 associated supplementary.**
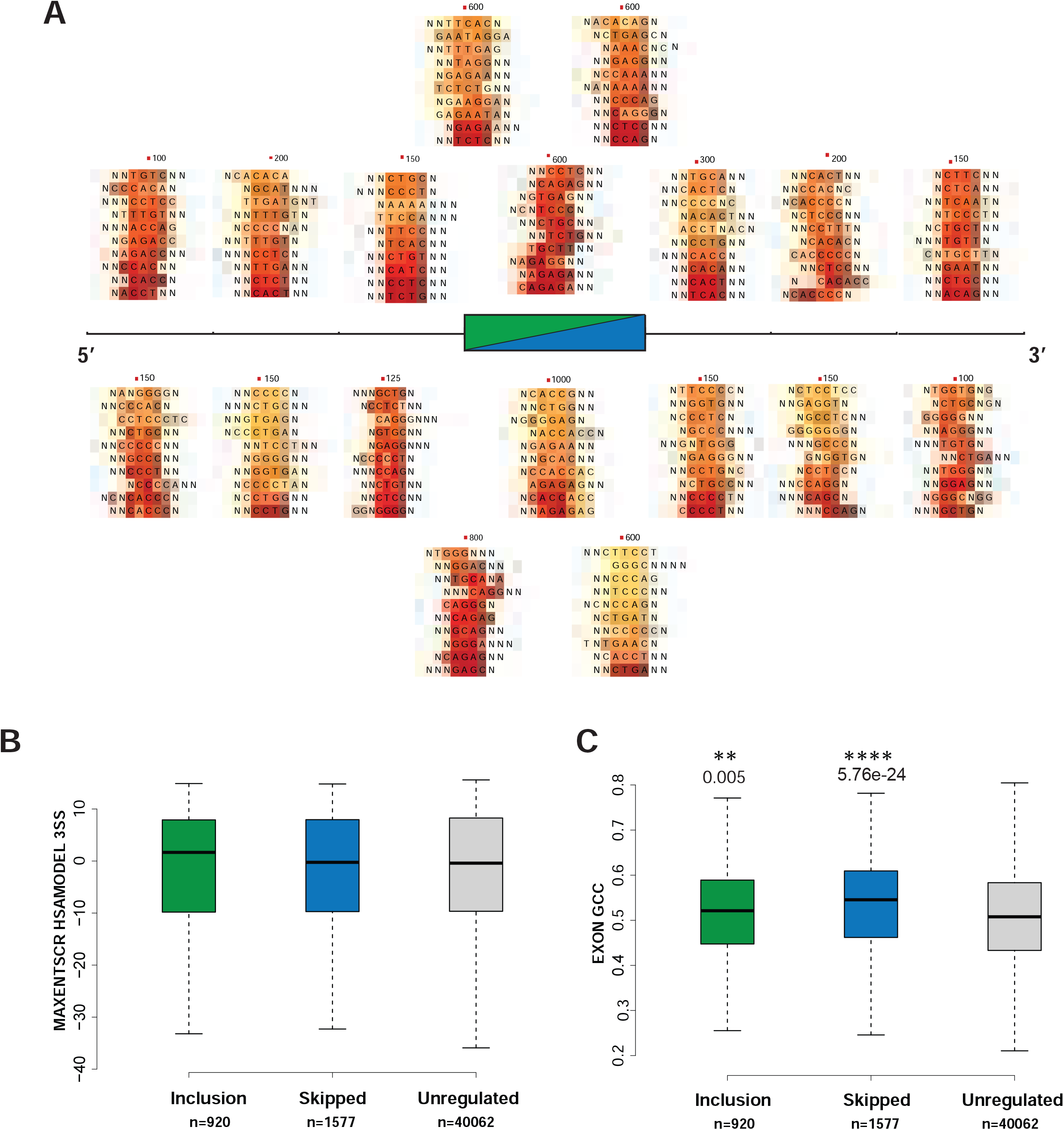
K-mer logo analysis shows enriched 5-mers on and flanking the RBPMS-regulated exons. **A**. 8-mer analysis using MATT of the RBPMS regulated exons (cassette exon category from rMATS) and 250bp of the flanking introns represented as motif logos. Test and background exons were designated as in Fig 5A. Heat maps depict enrichment scores of the various k-mers at each location on and flanking the regulated exons. **B**. RBPMS regulated exons do not show significantly different 3’ splice site strengths compared to the background exons. Analysis was with MATT suite using the MAXENT score model. Mann Whitney U test. **C**. RBPMS regulated exons have a significantly higher GC nucleotide content compared to the background exons. Mann Whitney U test.

**Fig 6 associated supplementary.**
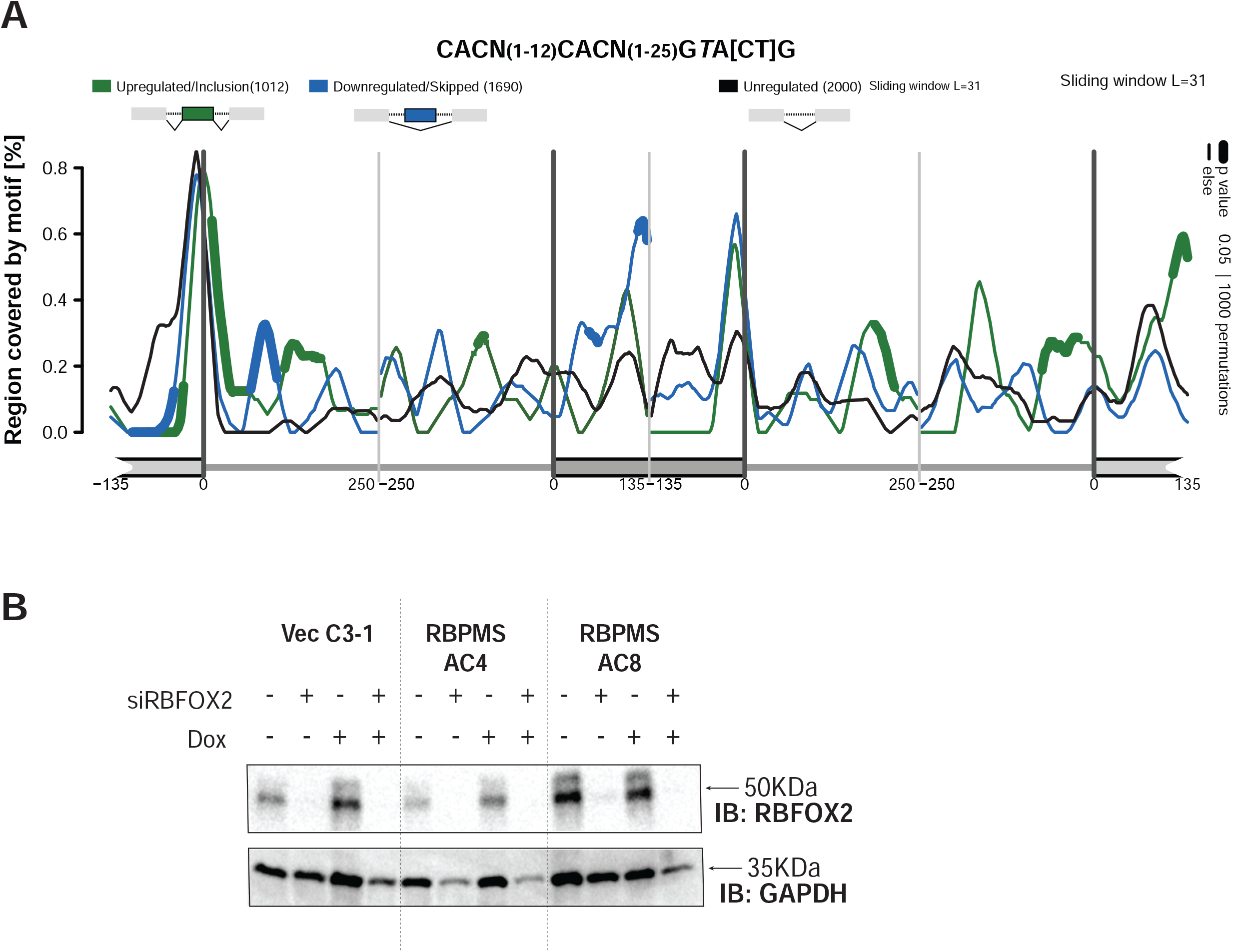
**A**. RNA map showing that disruption of the optimal RBFOX2 motif in the linked Perl expression results in a loss of enrichment suggesting that the co-occurrence of RBPMS binding motifs with RBFOX2 is specific. **B**. Immunoblotting showing knockdown of RBFOX2 in hES-VSMCs. GAPDH is used as a loading control.

**Fig 7 associated supplementary.**
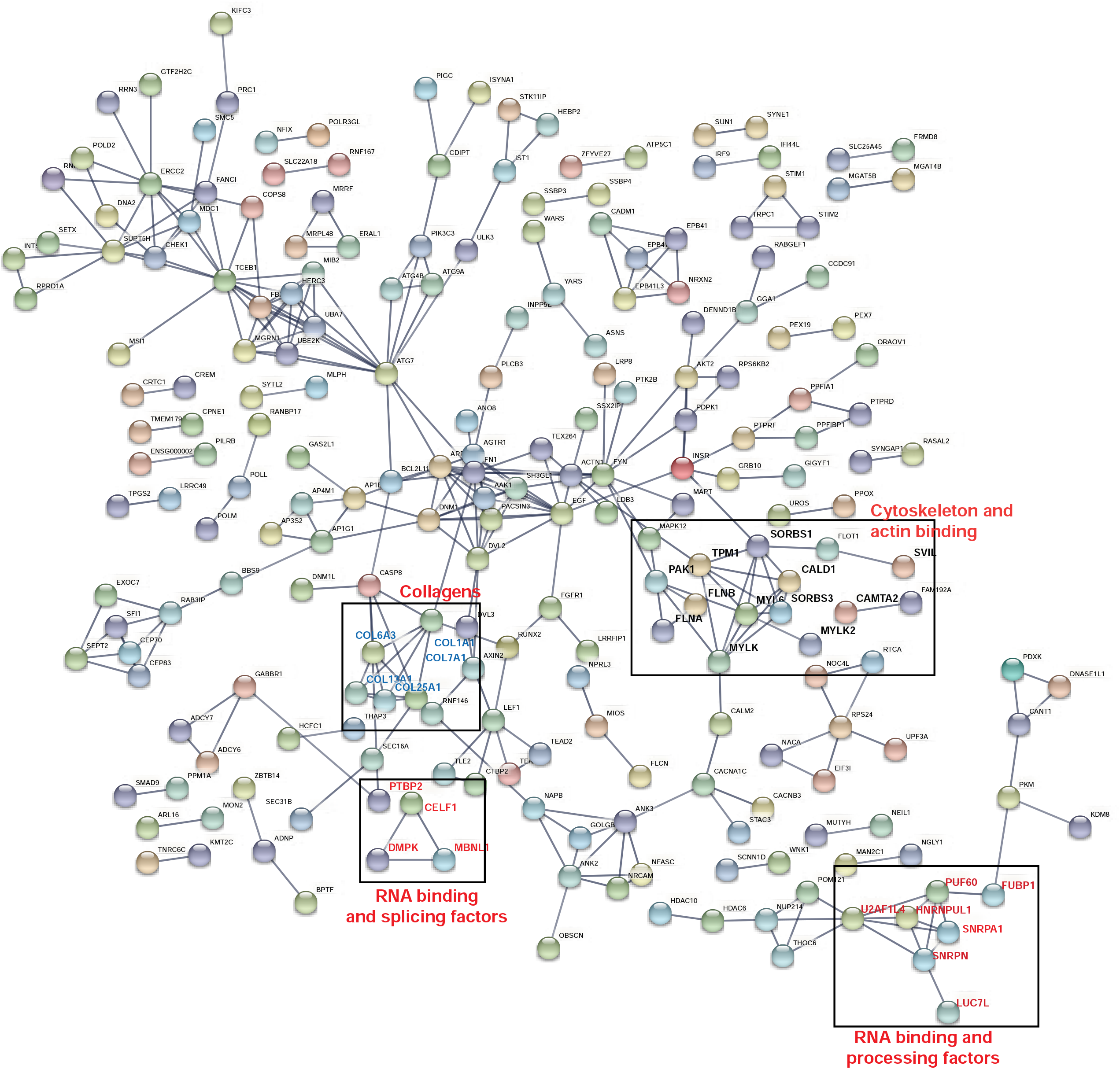
Protein-protein interaction networks of the RBPMS splicing network. Genes containing RBPMS regulated cassette exon events filtered for splicing difference 30% or greater and FDR < 0.05 were examined for protein-protein interaction networks using StringDB (string-db.org). 488 high confidence nodes with interaction scores of 0.75 or greater were determined connected by 219 edges. Background data was the whole human genome. Only query proteins are shown in the network represented here with no 1^st^ or 2^nd^ shell interactors depicted and disconnected nodes are not displayed. Gene clusters pertaining to cytoskeleton and actin machinery, RNA binding proteins and collagens are highlighted in bold within boxes.

**Fig 8 associated supplementary.**
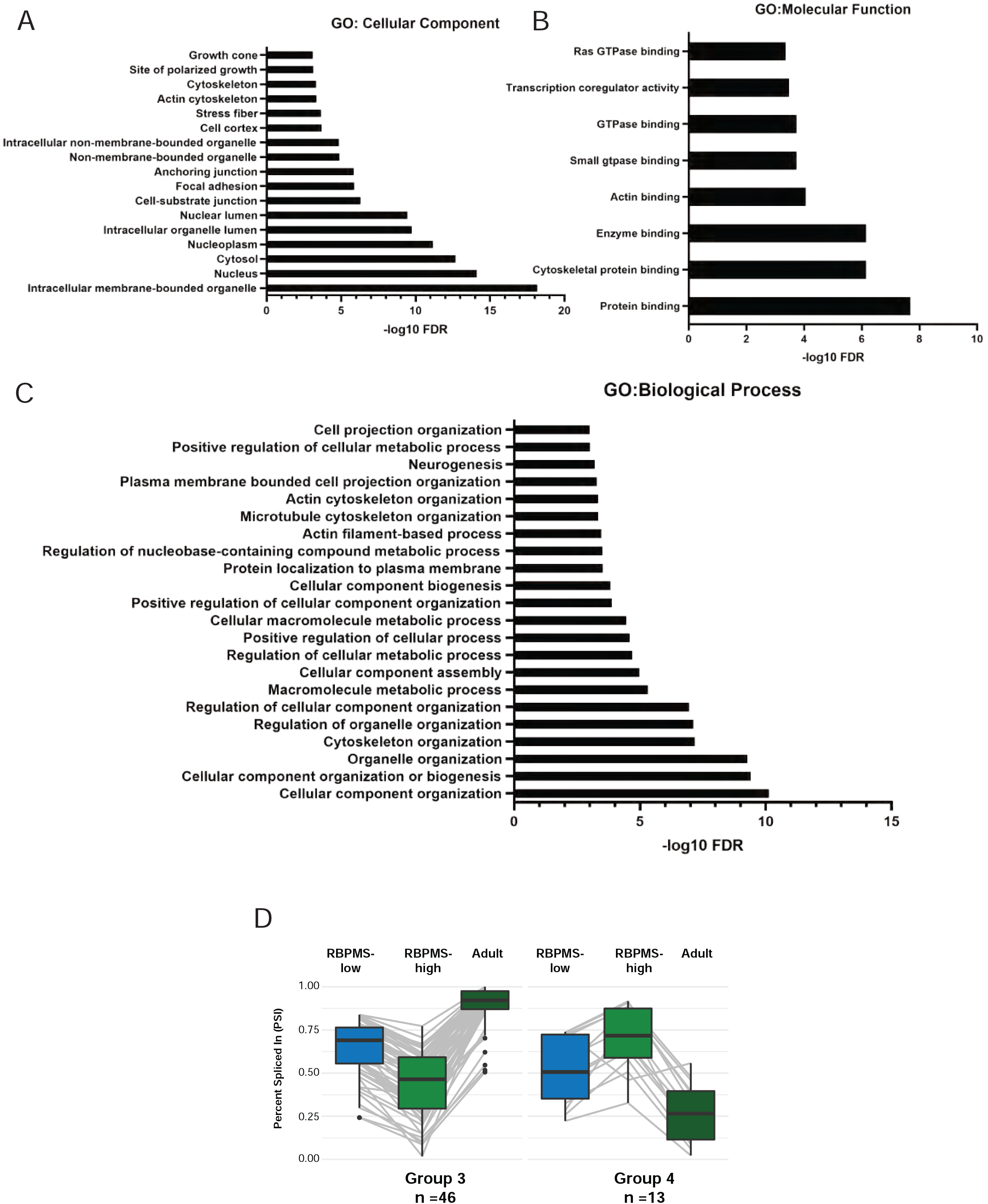
**A-C**. Genes showing significant alternative splicing (delPSI >=15% and FDR < 0.05) between adult human aorta and hES-VSMCs (RBPMS-low) representing the SM-AS events, were examined for protein-protein interaction networks with StringDB (string-db.org). Bar plots showing top enhanced enriched terms in the Molecular Function, Cellular Component and Biological Process categories respectively connecting high confidence nodes of interaction scores 0.99 or more. The terms enriched are represented with –log10 FDR and filtered from StringDB analyses for term strength (log 10 observed/expected) of 0.1 or more and FDR <= 0.001. SM-AS events are enriched for terms associated with the actin cytoskeleton and focal adhesion components (marked with arrow) similar to the RBPMS-regulated network indicating functional significance for VSMCs. **D**. Box plots showing the Percent Spliced In (PSI) distribution of the RBPMS-responsive non-SM-AS clusters 3 and 4 from the heatmap in 7A. In these events RBPMS regulation induces splicing patterns divergent from those seen in adult aorta tissue.

**Fig 9 associated supplementary.**
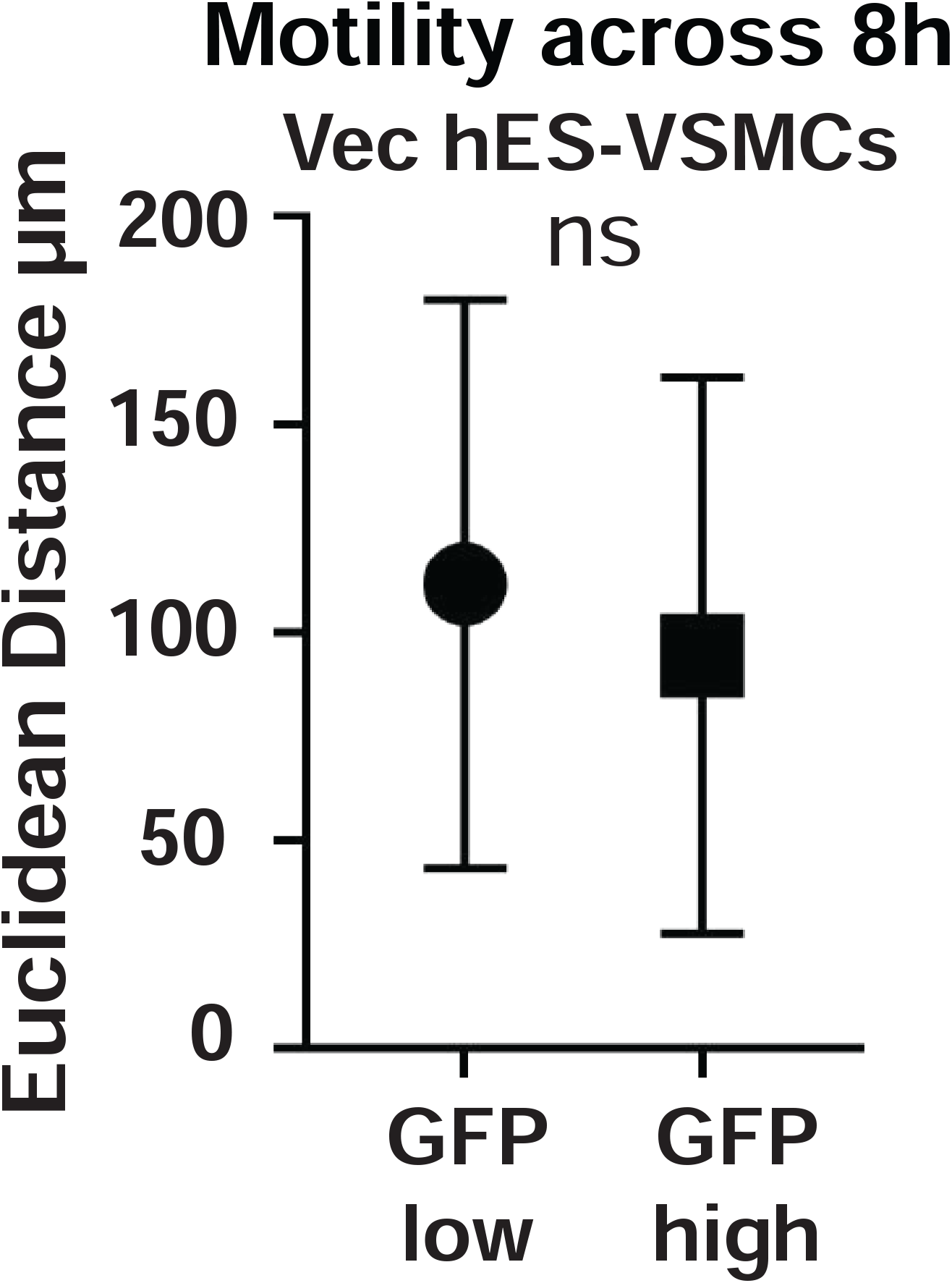
Motility of GFP high and low cells of 1 Vector control hES-VSMC line are shown. 40 cells across 2 wells were tracked in each category. GFP high and low cells do not show significant differences in their motility.

